# A single methylation site regulates HRSV nucleocapsid architecture and replication

**DOI:** 10.64898/2026.01.15.699754

**Authors:** Vincent Basse, Lorène Gonnin, Catarina S. Silva, Carine Rodrigues-Machado, Céline Henry, Maria Bacia-Verloop, Ambroise Desfosses, Julien Sourimant, Jean-François Eléouët, Cédric Leyrat, Irina Gutsche, Marie Galloux

## Abstract

The nucleoprotein N of the human respiratory syncytial virus (HRSV) encases the viral genome, forming a flexible N–RNA nucleocapsid helix that serves as template for the viral polymerase L. Recent structural analysis revealed a non-canonical helical nucleocapsid arrangement that modulates RNA accessibility, yet its impact on polymerase function remains unknown. Here, we identified symmetric dimethylation of residue R27 of N as a critical modulator of nucleocapsid architecture and viral replication. We also showed that the methylase PRMT5 interacts with N and likely catalyses R27 methylation. Molecular dynamics simulations of RNA-free N dimers indicate that R27 methylation enhances opening and closing of the RNA-binding cavity, whereas a methylation-mimicking R27M substitution favours a closed state. Cryo-electron microscopy reveals a canonical, markedly straight R27M helix with increased rise and pitch. These findings demonstrate that post-translational modifications fine-tune interactions between N protomers, shaping nucleocapsid assembly, structure and dynamics, and thereby controlling HRSV replication.

## INTRODUCTION

The human respiratory syncytial virus (HRSV), the prototype of the *Pneumoviridae* family within the *Mononegavirales* order ^1, 2^, is the leading cause of acute lower respiratory infections in young children worldwide. It leads to approximately 3.6 million child hospital admissions and 100,000 deaths annually, with the highest burden in developing countries ^3, 4^, and poses a major threat to the elderly and immunocompromised individuals ^5, 6^. The long-awaited first HRSV vaccines, Arexvy (GSK) and Abrysvo (Pfizer), were approved in 2023 but their use remains restricted to the elderly and to pregnant women ^7, 8^. Efforts to vaccinate children faced setbacks, with recent trials halted due to severe side effects ^9^. For now, the only available protection for infants consists in the injection of humanised monoclonal antibodies targeting the fusion protein F responsible for viral entry, such as Palivizumab (Synagis®) ^10, 11^ and Nirsevimab (Beyfortus®) ^12, 13^. While effective at preventing infection, these treatments offer only short-term protection, and HRSV-specific antivirals are still lacking.

HRSV possesses a non-segmented, negative-sense RNA genome ^12^, continuously and fully enwrapped by the viral nucleoprotein N. The resulting flexible helical nucleocapsid (NC) ^14, 15^ serves as the template for the viral RNA-dependent RNA polymerase L that transcribes and replicates the HRSV genome within cytoplasmic membraneless viral factories (VFs) ^16, 17^. Although encapsidation protects the viral genome from degradation by cellular RNases and from detection by innate immune sensors, it raises a long-standing question of how the L polymerase accesses the RNA to carry out its functions. A likely explanation is that NCs adopt distinct conformations, some permissive to polymerase activity and others not. Additionally, the transport of newly synthesised NCs from VFs to virion assembly sites at the plasma membrane might also depend on specific NC rearrangements and on interactions with cellular partners. The HRSV N protein consists of two globular domains (N_NTD_ and N_CTD_) connected by a hinge region that enables the inter-domain flexibility around the RNA binding groove ^18^. It also features N- and C-terminal extensions (NTD- and CTD-arms) that stabilise NC assembly. Notably, we recently uncovered a crucial role for the CTD-arm in driving the non-canonical helical organisation of recombinant HRSV NCs which results in periodic variations in RNA accessibility along the NC filament ^15^.

Another central player in HRSV genome transcription and replication is the P protein, a tetramer composed of a central oligomerisation domain flanked by mainly unfolded N- and C-terminal extensions, that recruits L to the NC. The interaction between NC and P occurs via binding of the C-terminus of P to the surface of N_NTD_ ^19, 20^, and is essential for VF formation ^21, 22^. In contrast, the N-terminus of P acts as a chaperone, stabilising newly synthesised N in a monomeric, RNA-free state (N^0^) and preventing its oligomerisation by anchoring to the surface of N_CTD_ ^23, 24, 25^. This mechanism ensures a readily available pool of N^0^ for selective encapsidation of HRSV antigenomes and genomes during replication. Recently, we demonstrated that phosphorylation of the residue Y88 of N stabilises the N^0^ form ^26^, underscoring the importance of post-translational modifications (PTMs) in regulating N structure and interactions, and suggesting that PTMs of monomeric N^0^ and oligomeric NCs might fine-tune transient protein-protein interactions required for the polymerase function on the NC template.

To explore this hypothesis and investigate potential roles of PTMs in modulating NC structure and L polymerase activity, we first searched for PTMs in HRSV NCs using mass spectrometry. We identified several PTMs in NCs isolated from infected cells and in recombinant NCs purified from insect cells. Among these, we chose to focus on the dimethylation of residue R27 of N, which was detected in both NC types. Combined biochemical and cellular assays revealed that mutations at R27 significantly affected both L activity and VF formation. The data further suggested that the cellular protein arginine methylase PRMT5 may be responsible for the symmetric dimethylation of R27. Using molecular dynamics (MD) simulations, we assessed the effects of such dimethylation and of a methylation-mimicking R27M mutation on the dynamics of N-N interactions, revealing a critical role of R27 in controlling N inter-domain motions relevant to RNA encapsidation. Finally, we leveraged cryo-electron microscopy (cryo-EM) to probe the structural impact of the R27M substitution on N-RNA assemblies produced upon expression of the R27M mutant of N in *E. coli*. Cryo-EM analysis of R27M NCs revealed a conformational rearrangement of the protomer, resulting in a remarkable shift in the NC polymorphism in comparison to the WT NCs and accompanied by a drastic reorganisation of the NC helical structure, which becomes canonical and rigid despite a dramatic increase in the helical rise and pitch. Together, our findings provide compelling evidence that PTMs, and in particular methylation, act as critical modulators of HRSV NC structure and function.

## RESULTS

### Identification of PTMs on HRSV nucleocapsids

To identify PTMs on HRSV N, HEp-2 cells were infected at high MOI with recombinant HRSV expressing fluorescent mCherry (rHRSV-mCherry), enabling infection efficiency to be validated via fluorescence microscopy ^27^. After 24 hours of infection, cells were lysed and N was immunoprecipitated for analysis by SDS-PAGE (Figure 1A). Coomassie blue staining showed a prominent band near the expected molecular weight of N (44 kDa). LC-MS/MS analysis confirmed that this band corresponded to HRSV N (99% sequence coverage, Table S1) and identified multiple N-derived peptides carrying PTMs, particularly within the NTD-arm. Considering that this region is crucial for N oligomerisation ^15, 18, 28^, we inspected the NTD-arm modifications more closely and were able to map them to phosphorylation at residues S20, S32, and Y38, as well as dimethylation at R27 (Table 1, Figure 1B). Notably, the peptide harbouring the R27 dimethylation yielded a particularly high confidence score (Table 1, Figure 1C).

**Figure 1:**
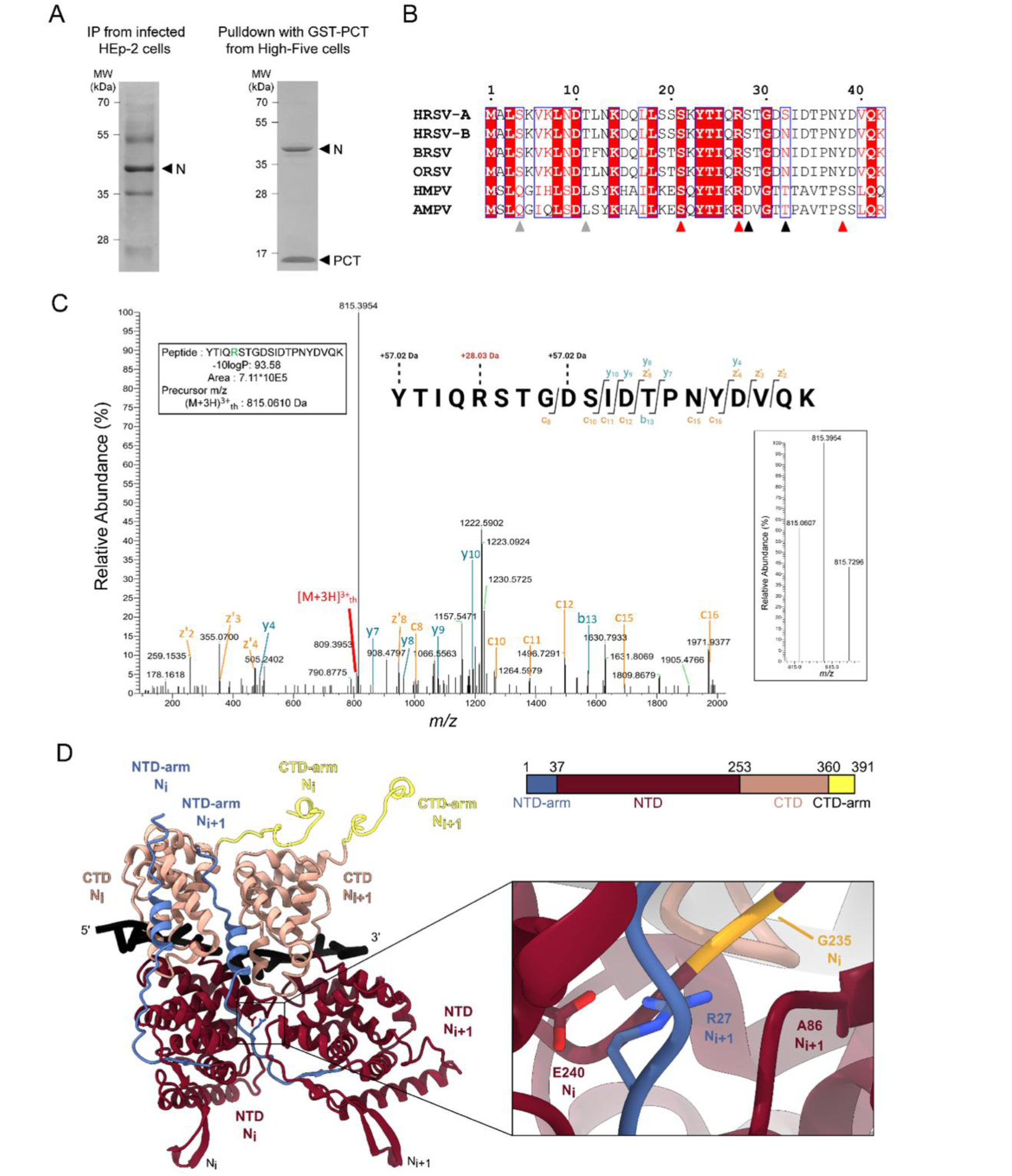
PTMs identification on HRSV nucleocapsids. **(A)** SDS-PAGE stained with Coomassie blue of N immunoprecipitated from infected HEp-2 cells using anti-N antibodies (left) or purified from High-five cells by affinity to GST-PCT (right). **(B)** Multiple sequence alignment of N sequences from *Pneumoviridae*. Invariant residues are highlighted in white font on a red background, and partially conserved residues are in red. Abbreviations and UniProt accession codes: HMPV (human metapneumovirus, NCAP_HMPVC), AMPV (avian metapneumovirus, NCAP_AMPV1), ORSV (ovine respiratory syncytial virus, NCAP_ORSVW), BRSV-A (bovine RSV type A, NCAP_BRSVA), MPV (murine pneumonia virus, NCAP_MPV15), HRSV-A (human RSV type A, NCAP_HRSVA), HRSV-B (human RSV type B, NCAP_HRSVB). Arrows below the sequences indicate the residues identified in both analyses (red), specific of N immunoprecipitated from infected cells (black), or of N purified from insect cells (grey). **(C)** LC-MS/MS fragment coverage map for the assignment of R27 dimethylation on the YTIQRSTGDSIDTPNYDVQK peptide, identified for the N protein immunoprecipitated from infected HEp-2 cells, and digested with trypsin. The overall observed mass difference to three residues is indicated: Y1 (+57.02 Da), R5 (+28.03 Da) and D9 (+57.02 Da). A mass gain of 57.02 Da is typical of a carbamido-methylation event that likely occurred during the MS analysis through reaction with iodoacetamide, while +28.03 Da indicates a dimethylation event. MS/MS data is shown with z’- and c-ions depicted in orange, and y- and b-ions in cyan (see Table S1). The isolation window is shown on the right. **(D)** Atomic model of two consecutive protomers from HRSV WT N_10_ double rings (PDB: 8OOU), shown as ribbons. The protomers are coloured as per the schematic of the HRSV N sequence ^15^ shown, with the NTD-arm in blue-grey, the NTD in rosewood, the CTD in old rose, and the CTD-arm in yellow. The RNA is in black. The close-up shows the interactions between the residue R27 of the protomer N_i+1_ with the residues E240 and G235 of the protomer N_i_, as well as A86 from the protomer N_i +1_.

**Table 1:**
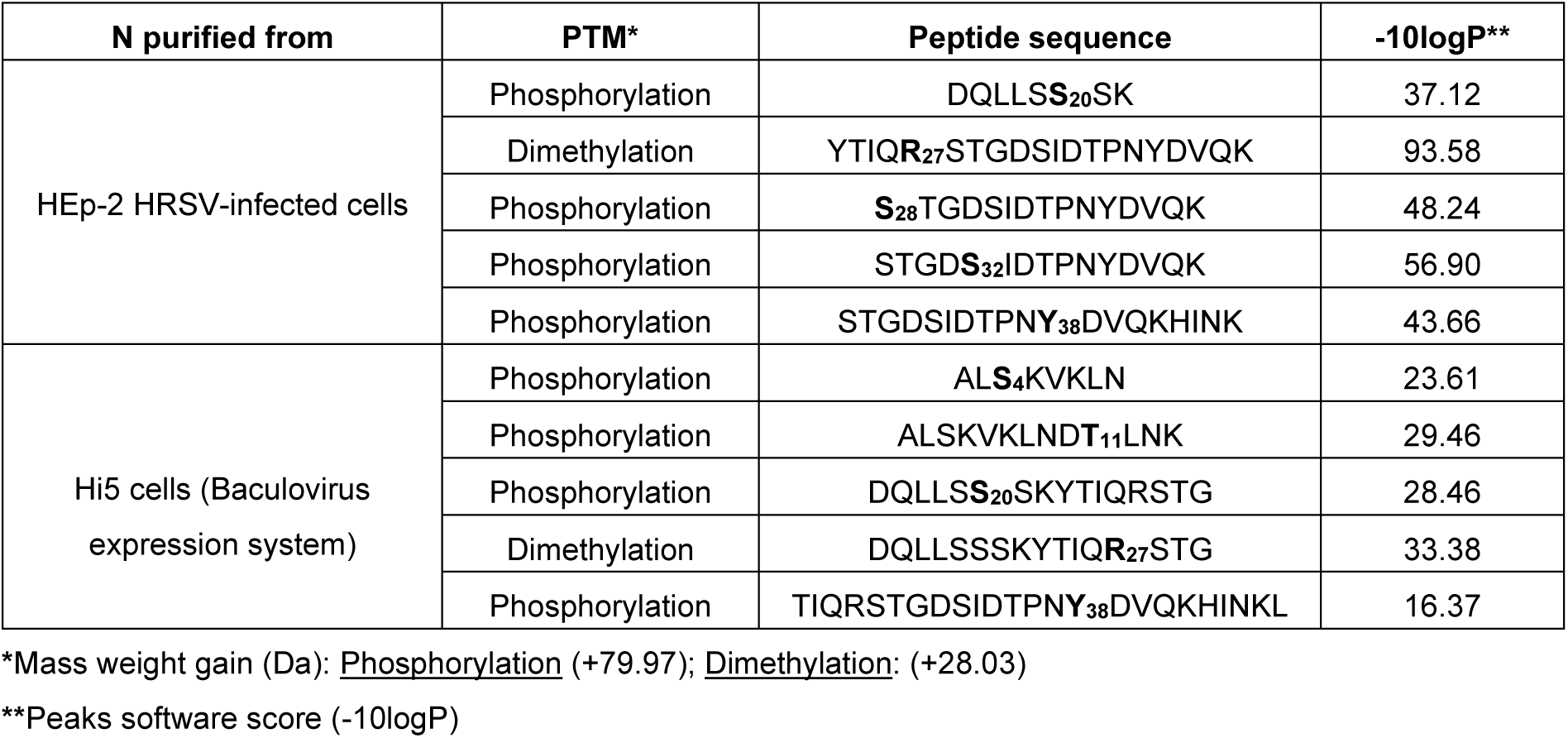
Modified peptides identified by LC-MS/MS.

Since immunoprecipitation does not allow to distinguish between N^0^ and oligomeric N-RNA, we next examined PTMs in recombinant HRSV NCs purified from insect cells. Using the baculovirus expression vector system, HRSV NCs were produced in High-Five cells and purified via affinity chromatography with a GST-fused C-terminal domain of P (GST-PCT), as previously described ^15^ (Figure 1A). In contrast to N immunoprecipitated from infected cells, SDS-PAGE of these purified NCs showed only two clear bands corresponding to N and GST-PCT. LC-MS/MS analysis of the N band detected phosphorylation at S4, T11, S20, and Y38, alongside a dimethylation at R27 (Table S1). Again, the R27 dimethylated peptide displayed the highest confidence score. Thus, our observations demonstrate that phosphorylation at S20 and Y38, along with dimethylation at R27, occur in both N immunoprecipitated from infected cells and recombinant NCs produced in insect cells. Notably, Y38 phosphorylation was previously shown to be essential for HRSV polymerase activity ^29^, suggesting that these PTMs may play a role in the regulation of N structure and function. Additionally, our previous work identified PTMs in the NTD-arm of N that are involved in stabilising the N^0^ form ^26^. However, none of these modifications were detected in the present analysis, implying that PTMs may be specific to certain oligomeric states and conformations of N.

Multiple sequence alignment of *Pneumoviridae* N proteins revealed that among the identified modified residues, R27 is the only one conserved (Figure 1B). In the NC structures, R27 of the protomer N_i+1_ is involved in key interactions with A86 of the same protomer, and with G235 and E240 of the protomer N_i_ (Figure 1D). Notably, previous studies have shown that mutating this residue in the N protein of human metapneumovirus (HMPV) impairs L polymerase activity ^30^. Altogether, these findings indicate that HRSV NCs can undergo various PTMs, with R27 dimethylation standing out as a potential regulator of N oligomerisation and, by extension, polymerase activity.

### R27 mutations strongly affect viral polymerase activity and VF formation

To assess the impact of R27 mutations on viral polymerase activity, we employed the minigenome assay, as previously described ^20, 31^. R27 was substituted with alanine, lysine, or methionine — mutations expected to disrupt lateral chain interactions at this position by altering charge and steric hindrance (R27A, R27K), or mimicking methylation (R27M). As a control, we also examined the effect of mutating Y38 to phenylalanine (Y38F), which preserves the aromatic side chain, or to aspartic acid (Y38D), which mimics phosphorylation. Strikingly, both R27A and R27M completely abolished L polymerase activity, while R27K retained only residual L activity of approximately 10% of the WT level (Figure 2A). Conversely, Y38F had no impact on polymerase function, whereas Y38D nearly eliminated L activity, in agreement with previous findings ^29^. Importantly, none of these mutations affected N expression, as confirmed by western blot analysis (Figure 2B).

**Figure 2:**
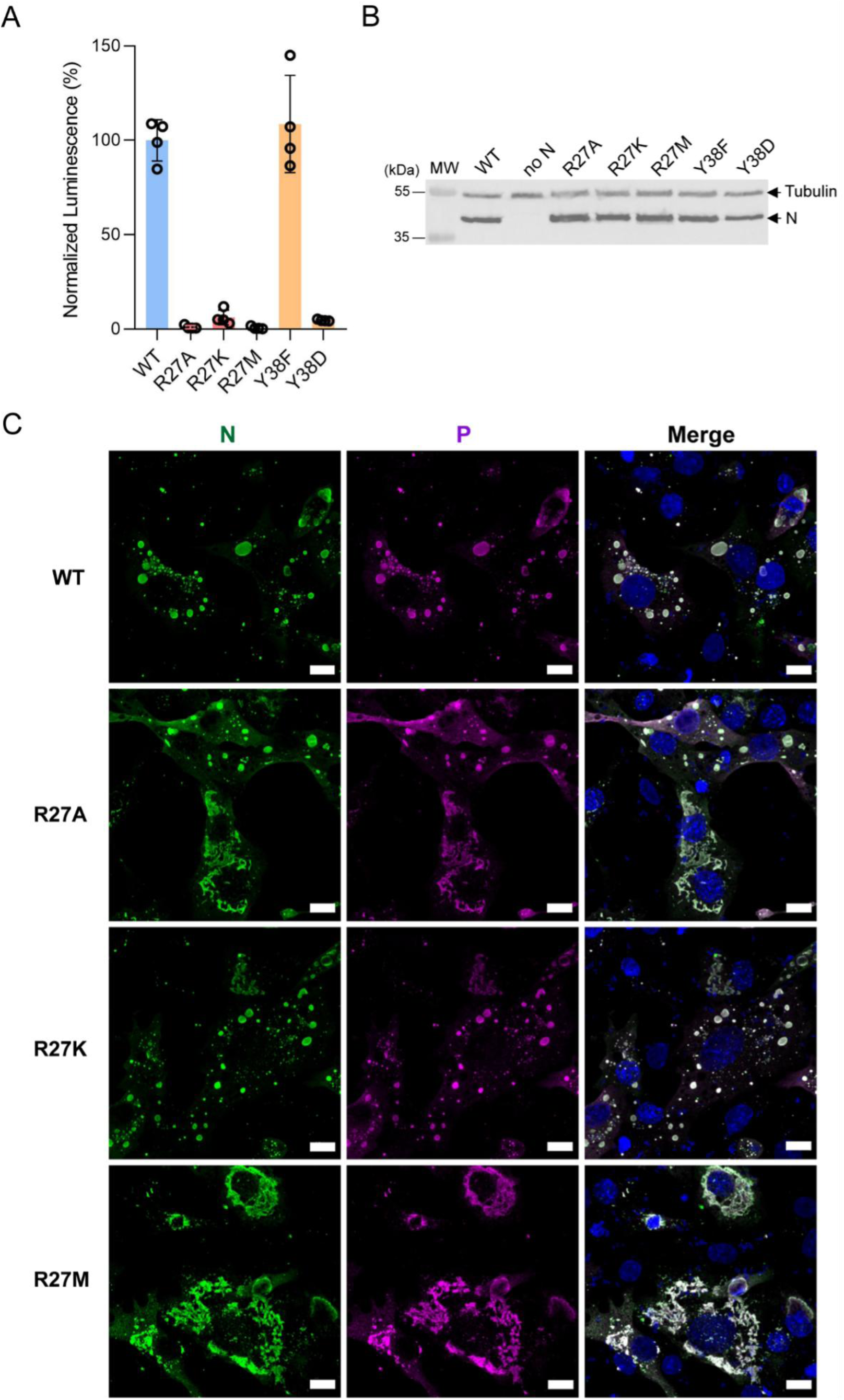
Impact of R27 substitutions on HRSV polymerase activity and pseudo-VFs formation. **(A)** BSRT7/5 cells were transfected with plasmids encoding the WT P, M2-1 and L proteins, the pMT/Luc minigenome, and WT or R27 and Y38 mutants of the N protein, together with pCMV-βGal for transfection standardization. Cells were lysed 24 h post-transfection and viral RNA synthesis was quantified by measuring the luciferase activity. Each luciferase minigenome activity value was normalized based on β-galactosidase expression. The experiment was performed three times. Data points represent values of quadruplicates from one representative experiment. Error bars represent standard deviations (+/-SD). **(B)** Western blot showing the expression of the N protein (WT and mutants) and tubulin as a control, in BSRT7/5 cells**. (C)** BSRT7/5 cells were transfected with plasmids encoding N (WT or R27 mutants) and P, fixed 24 h post-transfection, and labeled with anti-N (green) and anti-P (red) antibodies. The localisation of N and P proteins was observed by fluorescence microscopy. Nuclei were stained with Hoechst 33342. Scale bars, 10 µm.

Next, we investigated whether R27 mutations influenced the cellular localisation of N, or more specifically, its ability to form pseudo-VFs when co-expressed with P ^16, 17, 32^. To this end, cells were co-transfected with plasmids encoding P and either WT or mutant N, and their localisation was examined by immunolabelling and subsequent confocal microscopy (Figure 2C). While R27K allowed the formation of pseudo-VFs similar to those observed with WT N, the substitutions R27A and R27M significantly disrupted the process. Interestingly, R27A exhibited two distinct phenotypes: N and P either formed spherical cytoplasmic inclusions resembling pseudo-VFs or displayed diffuse cytoplasmic aggregates. In contrast, R27M led exclusively to the formation of diffuse cytoplasmic aggregates where N and P colocalised (Figure 2C). These findings suggest that R27 mutations may induce conformational rearrangements of HRSV NCs that impair the ability of N and P to undergo liquid-liquid phase separation, the R27M mutation exerting the most profound effect. Taken together, these results further highlight the crucial role of R27 in HRSV polymerase activity and suggest that its methylation could play a key role in NC organisation.

### Characterisation of the symmetric nature of R27 dimethylation

We then sought to determine the nature of R27 dimethylation. Arginine methylation depends on protein arginine methyltransferases (PRMTs), which catalyse transfer of methyl groups from the co-substrate S-adenosylmethionine (SAM) to nitrogen atoms in the guanidino group of arginine ^33^ (Figure S1). The process begins with the addition of a single methyl group to one of the terminal nitrogen atoms of the arginine residue, resulting in mono-methylation (MMA). A second methylation step can then produce either asymmetric dimethylation of the arginine (ADMA), when performed by type I PRMTs (PRMT1, 2, 3, 4, 6, 8), or symmetric dimethylation of the arginine (SDMA), catalysed by type II PRMTs (PRMT5, 9). While most PRMTs preferentially target arginine residues within RG or RGG (Arg-Gly-Gly) motifs, they can also methylate arginine within non-specific sequences. Deciphering the nature of R27 dimethylation is challenging due to limited available tools. While commercial antibodies exist for detecting ADMA (anti-adme-R antibodies), SDMA-specific antibodies are typically designed for RG motifs (anti-sdme-RG antibodies) or require multiple SDMA sites for detection. Therefore, potential SDMA on R27, which is not located within an RG motif, cannot be detected with these antibodies. To assess ADMA on N, we analysed both NCs purified from lysates of either infected HEp-2 cells or of insect cells expressing N (Figures 3A and S1). The anti-adme-R antibody detected multiple bands in lysates of both non-infected and infected HEp-2 cells, but no differences were observed between conditions, and most importantly, no signal corresponding to N was detected, as confirmed by anti-N antibody staining (Figure 3A). Similarly, in lysates of insect cells, the anti-adme-R antibody revealed several bands, and predominantly one that could correspond to the molecular weight of N. However, no signal was detected in the sample of purified NCs, despite N presence being confirmed using an anti-N antibody (Figure S1). While we cannot entirely rule out the presence of ADMA on HRSV N that may have escaped detection, our results suggest that R27 is more likely the site of SDMA. Of particular interest, PRMT5, the predominant and ubiquitously expressed SDMA-catalysing type II PRMT ^34^, was previously found to co-purify with HRSV N expressed in 293T cells ^35^, and was recently shown to methylate the PB2 subunit of the influenza polymerase ^36^.

**Figure 3:**
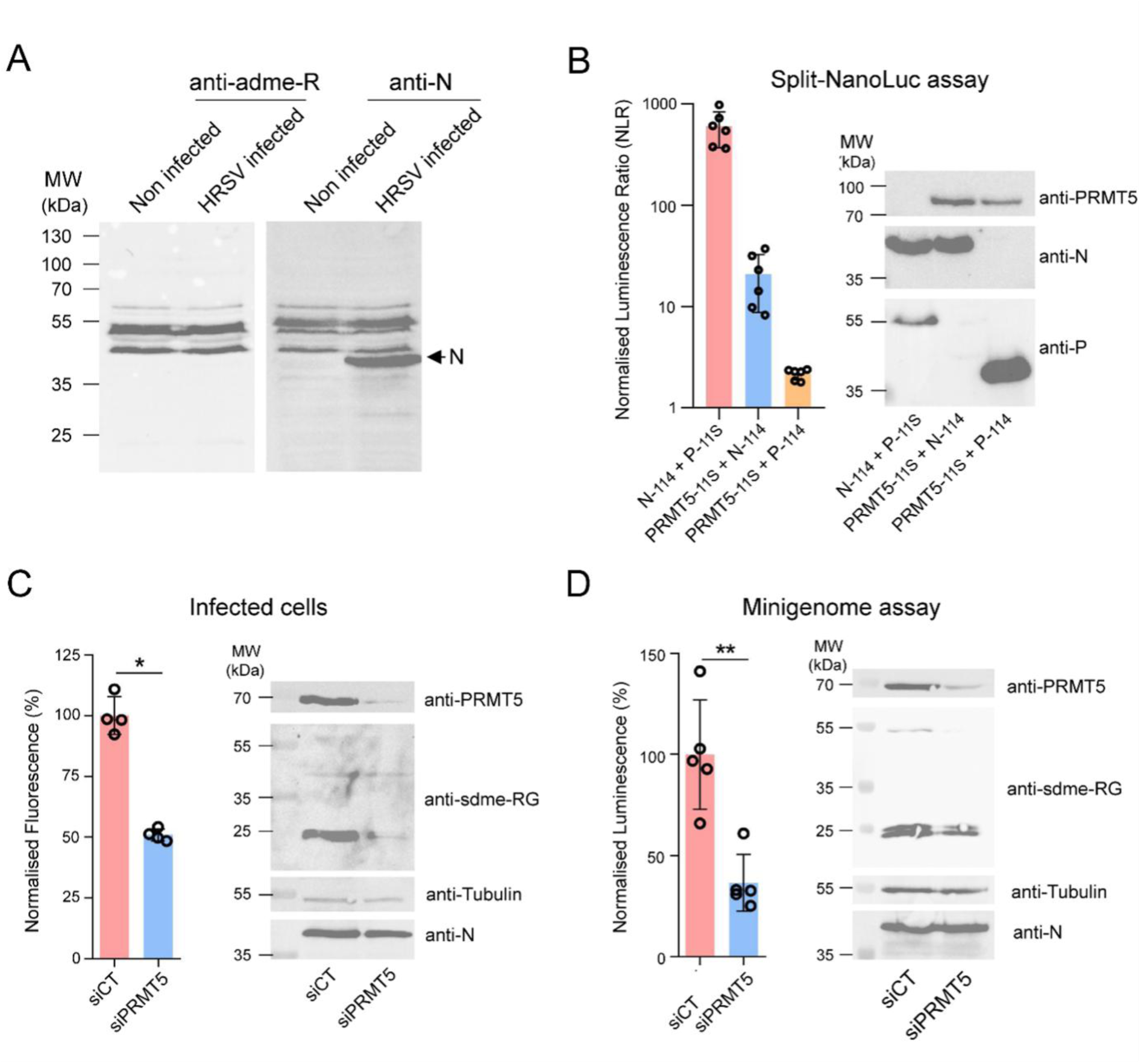
Investigation of the potential role of PRMT5 in N methylation. **(A)** Western blot analysis of the profile of ADMA proteins in the lysates of non-infected or infected HEp-2 cells. HEp-2 cells were infected with rHRSV-mCherry at MOI 1 for 24 h. **(B)** Study of N-PRMT5 interaction using the split NanoLuc assay. HEK293T cells were transfected with pairs of plasmids encoding either N-114 with P-11S or PRMT5-11S, or with PRMT5-11S and P-114. Protein-protein interactions were measured by quantification of the luminescence. The NLR, which corresponds to the ratio between actual read and negative controls (each protein with the complementary empty NanoLuc vector), was determined. The graph is representative of four independent experiments, each done in six technical repeats. Error bars represent standard deviations (+/- SD) calculated based on one experiment (left). Protein’ expression was validated by western blot analysis (right). **(C)** Impact of PRMT5 depletion on HRSV replication in cells. A549 cells were transfected with control siRNA (siCT) or siRNA targeting PRMT5 (siPRMT5) and infected 24 h later with rHRSV-mCherry at MOI of 0.2. HRSV replication was quantified at 48 h post-infection by measurement of fluorescence (left), in cell culture. Data are representative of results from three experiments made in quadruplicates. Error bars represent standard deviations (+/- SD) calculated based on one representative experiment. *, *P* < 0.05. Western blot analysis of PRMT5 silencing, SDMA level of proteins, HRSV N and tubulin expression in infected cells at 48 h post-infection (right). **(D)** BSRT7/5 cells were transfected with control siCT or siPRMT5 and transfected the day after with the plasmids of the minigenome assay. Cells were lysed 24 h later and the polymerase activity was quantified by measuring the luciferase activity (left). The experiment was performed four times and a representative experiment done in five technical repeats. Each luciferase minigenome activity value was normalized based on β-galactosidase expression and is the average of quadruplicates. Error bars represent standard deviations (+/- SD) calculated based on one representative experiment. **, *P* < 0.01. Western blot analysis of PRMT5 silencing, SDMA level of proteins, HRSV N and tubulin expression in cells (right).

### PRMT5 involvement in N binding and regulation of HRSV polymerase activity

To address the potential role of PRMT5 in N methylation, we first sought to confirm the interaction between N and PRMT5 ^35^. For this, we employed a split-luciferase complementation assay based on the Nano Luciferase (NanoLuc) enzyme ^37^. In this assay, the 114 peptide or the 11S large domain of the NanoLuc were fused to the C-terminus of each protein partner. Pairs of constructs were transfected into 293T cells, which were lysed 24 hours post-transfection, and the luminescence intensity, proportional to the strength of interaction, was then quantified following substrate addition. The interaction between HRSV N and P proteins was used as a positive control. As shown in Figure 3B, co-transfection of N-114 and P-11S resulted in a high normalised luminescence ratio (NLR, close to 600), confirming the N-P interaction in this system. When testing the interaction between PRMT5-11S and either N-114 or P-114, our results revealed an interaction between N and PRMT5 (NLR close to 20), while no relevant interaction was detected between P and PRMT5 (NLR < 3). The expression of the fusion proteins was confirmed by western blot (Figure 3B).

Next, we assessed the impact of PRMT5 downregulation on HRSV replication. For this experiment, we used human epithelial A549 cells, which are more permissive to siRNA transfection than HEp-2 cells. Cells were transfected with siRNA for 24 hours, then infected with rHRSV-mCherry at an MOI of 0.2 for 48 hours. Fluorescence-based quantification of viral replication showed that PRMT5 downregulation led to a 50% decrease in infection (Figure 3C). Western blot analysis confirmed reduced PRMT5 expression, accompanied by a decrease in global SDMA signal in the cell lysate (Figure 3C), while N expression was only mildly affected. We further examined the effect of PRMT5 silencing on polymerase activity using the minigenome assay. As this assay is performed in BSRT7/5 cells, a hamster cell line, we first verified the sequence conservation between human and hamster PRMT5, which turned out to be extremely high (98%), suggesting that the human siRNA would likely function in this system also. Cells were transfected with either siCT or siPRMT5 for 24h before transfection with the minigenome system. Once again, PRMT5 downregulation resulted in decreased L polymerase activity (Figure 3D). Western blot analysis confirmed the efficiency of siPRMT5 transfection, showing reduced PRMT5 expression and lower SDMA levels in the cell lysate (Figure 3D). Notably, siRNA treatment did not affect N protein production, as N expression in this assay is controlled by the T7 promoter and mediated by T7 polymerase expressed in the cytoplasm of BSRT7/5 cells.

We next sought to assess the potential recruitment of PRMT5 into pseudo-VFs formed upon co-expression of N and P, as well as into VFs of HRSV infected cells, using immunolabelling to detect endogenous PRMT5. The available commercial antibodies, however, did not yield a detectable signal. To overcome this limitation, we also overexpressed tagged PRMT5, but its high expression levels proved toxic to cells, precluding reliable analysis of potential colocalisation of PRMT5 with N.

In summary, although our results did not provide definitive evidence that PRMT5 methylates N, they confirm that PRMT5 can interact with N and plays a crucial role in the HRSV polymerase activity.

### MD simulations highlight the impact of R27 modification on the dynamics of N–N interactions

In order to gain a detailed atomistic understanding of the role of R27 in N-N interactions, we performed MD simulations of an RNA-free N_i_-N_i+1_ dimer in which R27 was either kept as a regular arginine residue, dimethylated (dmR27) or substituted with a methionine (R27M) (Figure 4A-C). Although simplified, this system reproduces the N-N interface, including the N_i+1_ NTD-arm sandwiched between the two protomers. In addition, the absence of bound RNA allows to monitor the intrinsic dynamics of N, relevant to RNA encapsidation. As mentioned above (Figure 1D), in the case of unmodified R27 N dimers, R27 side chain of the protomer N_i+1_ maintains a stable network of polar contacts during MD, involving N_i+1_ A86 backbone carbonyl, and N_i_ R234 and G235 backbone carbonyl, as well as the E240 carboxylate (Figure 4A and S2A). In contrast, the presence of 2 methyls on dmR27 guanidinium group leads to a loss of hydrogen bonding with A86 and R234 and increased local flexibility (Figure 4B and S2B). However, dmR27 still maintains an ionic interaction with E240 as well as other dynamic, relatively short-lived contacts. The R27M mutant also loses the hydrogen bonding possibilities offered by the arginine guanidinium group, its shorter sidechain leaving a water-filled void in this part of the interface (Figure 4C and S2C). Taken together, these results indicate that both dimethylation and R27M substitution strongly impact N-N_i_ interactions, with an important increase in side chain flexibility observed for dmR27. To determine the effect of N_i+1_ R27 modifications on the large-scale conformational dynamics of the N_i_ protomer interacting with modified N_i+1_ NTD-arm, we applied principal components analysis (PCA) to the N_i_ protomer core extracted from MD trajectories of WT, dmR27 and R27M RNA-free dimer systems (Figure 4D-I). The analysis showed that N dynamics is dominated by concerted inter-domain motions corresponding to the opening/closure of the RNA binding cavity separating the NTD and CTD (PC1), and the relative twisting of the NTD and CTD (PC2) (Figure 4D-E). These motions are typical for *Mononegavirales* N proteins due to their conserved shape, and similar PCs have been described in previous studies of rabies virus N protein ^38^. The PC1 opening/closure and PC2 twisting motions capture roughly 60% of the variance observed in the MD dataset (Figure 4F). Two-dimensional projections of the MD trajectories along PC1 and PC2 for each system reveal important differences in dynamics (Figure 4G-I). In the case of the native R27 system, the N_i_ protomer samples various conformational states around the starting experimental structure characterised by moderate opening/closure (PC1) and important twisting motions (PC2) (Figure 4G). In contrast, dmR27 shows a comparatively flattened profile along PC2, and an enhancement of opening/closure motions along PC1 leading to an equilibrium with more open conformations not sampled by the native R27 system (Figure 4H). Finally, R27M shows a clustering of the structures in a region of PC space corresponding to closed conformations (Figure 4I), indicating that the loss of polar interactions combined with the shorter methionine sidechain in N_i+1_ NTD-arm increase the relative stability of the closed state of N_i_ protomer. Altogether, the MD simulations identify R27 as central to the dynamics of N–N interactions, which govern the large-scale N opening motions essential for RNA encapsidation.

**Figure 4:**
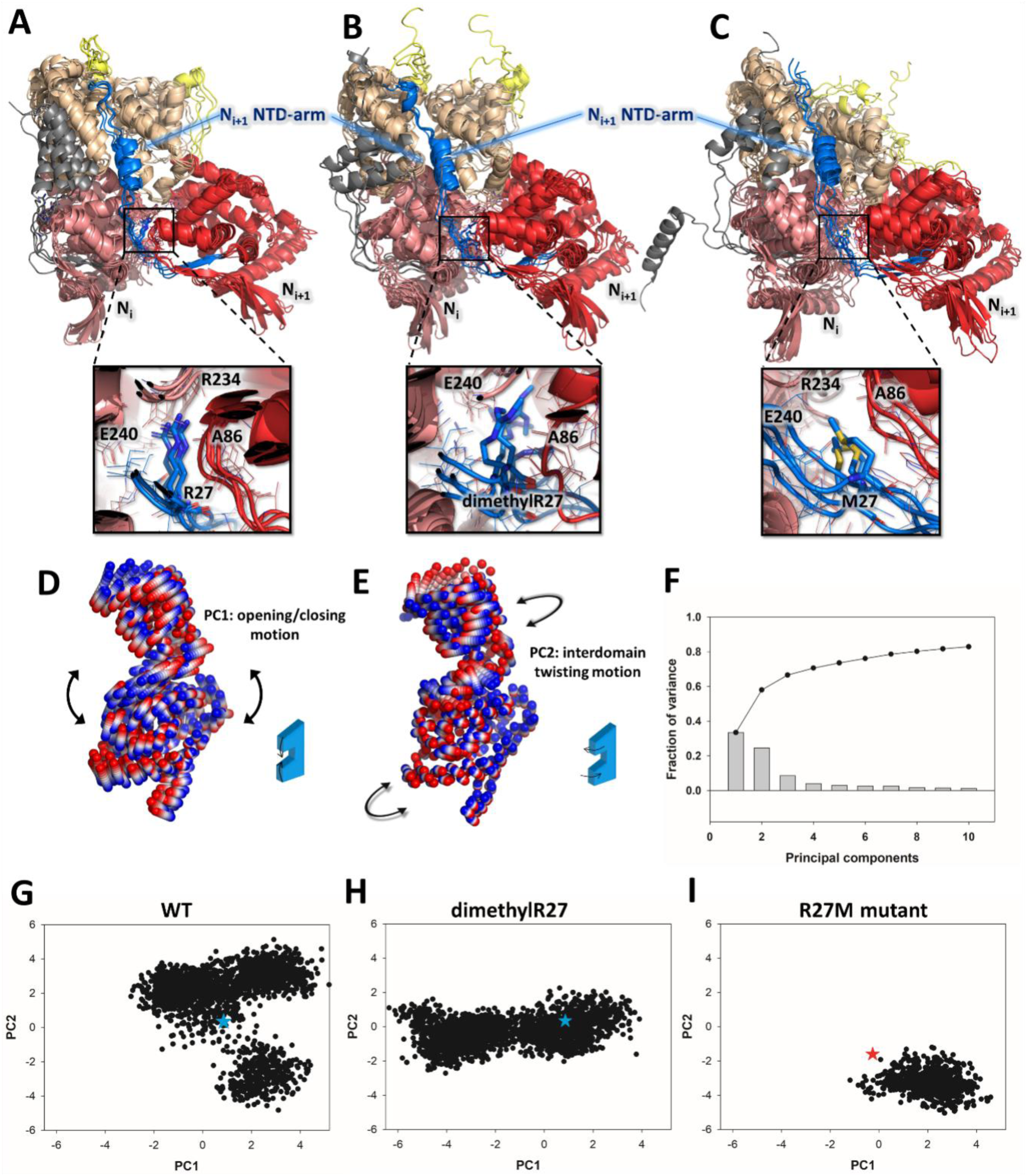
Effect of R27 symmetric dimethylation and R27M substitution on N-N interaction dynamics studied by MD simulations. **(A)**, **(B)** and **(C)**. Snapshots of the N-N dimers extracted from representative MD trajectories of unmodified R27 (A), dmR27 (B) and R27M (C) systems. The snapshots were taken at t=0, 200, 400 and 600 ns. The N_i+1_ NTD is coloured in red, with its NTD-arm bearing the (modified) R27 residue in blue. The N_i_ NTD is coloured in salmon with its NTD-arm in dark grey. The CTD and CTD-arms of both protomers are coloured in light brown and yellow, respectively. The insets show close up views of unmodified R27, dmR27 and R27M in the MD simulation snapshots. **(D)** to **(I)**. Principal component analysis (PCA) of the N_i_ protomer extracted from MD simulations. **(D)** and **(E)**. Collective motions of N captured by PC1 and PC2. The motions are illustrated as linear interpolations between the extreme projections of the structures onto the PCs. Each cylinder thus describes the path of a Cα atom between its extreme positions (on a blue–white–red color scale). **(F)**. Fraction of the variance captured by each PC (histogram) and cumulative contributions of the first ten PCs. **(G)**, **(H)** and **(I)**. 2D projections of the first two principal components (PCs) calculated for the N_i_ protomer from MD simulations of N-N dimers with unmodified R27 (G), dmR27 (H) and R27M (I). The values calculated from the starting experimental structures are shown as a blue (PDB 8op2, WT) and a red star (R27M structure).

### Recombinant R27M NCs display a unique polymorphism, helical canonicity and a prominent rise

In parallel to exploring the role of R27 in an RNA-free N_i_-N_i+1_ dimer by MD simulations, we expressed the R27M-N mutant in *E. coli* (Figure S3) to assess the impact of this methyl-mimetic mutation on N oligomerisation. Notably, although the purified R27M-N did contain RNA (OD_260_/OD_280_ ratio of 1.5), its migration profile on both native agarose or acrylamide gels differed from that of the WT (Figure S3). Cryo-EM observation of the R27M-N preparation immediately explained this difference: instead of expected 10 and 11-fold symmetric N-RNA rings, typically observed for WT HRSV N produced in *E. coli* ^18^ and separating into two bands on a native acrylamide gel (Figure S3C), purified R27M N-RNA displayed short helical spirals and even NC-like filaments (Figure 5A). Noteworthy, the WT N of HMPV, closely akin to HRSV but belonging to the *Metapneumovirus* rather than the *Orthopneumovirus* genus, was also shown to assemble into spiral particles when produced in *E. coli*, although, in contrast to our HRSV R27M preparation, the usual 10 and 11-mer N-RNA rings were predominant ^30^. The high-resolution structures of the HRSV R27M N-RNA spirals and helices are presented and discussed in detail below.

**Figure 5:**
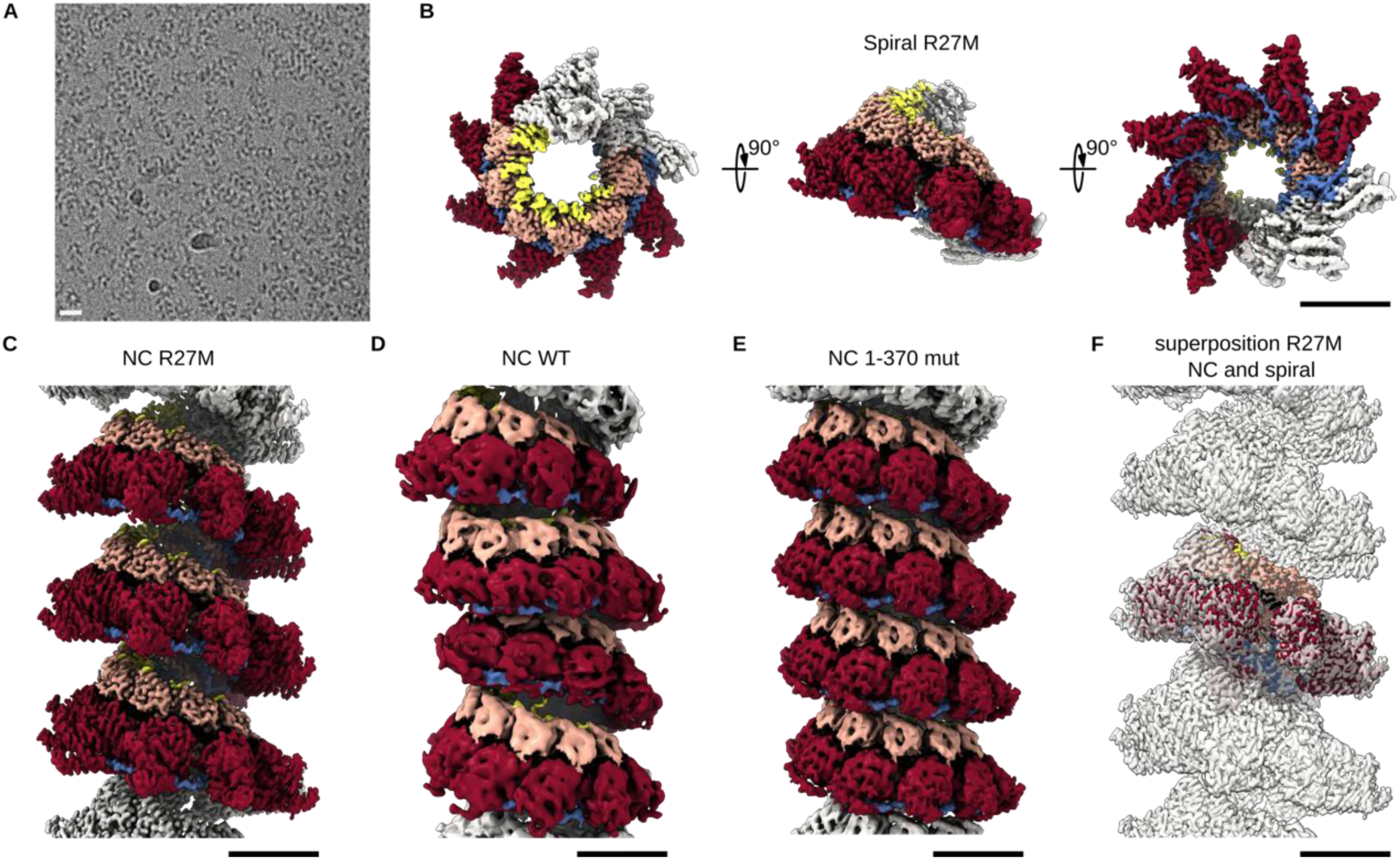
Cryo-EM analysis of the HRSV R27M NC helices and spirals. Colouring as in Figure 1D. **(A)** Representative micrograph of HRSV R27M NCs purified from *E. coli*, featuring spiral and filamentous NCs, scale bar 200 Å. 20,495 micrographs were selected for further processing. **(B)**. Cryo-EM map of the R27M NC spiral (EMD:56158), top, side and bottom view. **(C)** Cryo-EM map of the R27M NC helix (EMD:56157). **(D)** Cryo-EM map of the non-canonical WT NC helix purified from insect cells (EMD:17030). **(E)** Cryo-EM map of the canonical N1-370 NC helix purified from insect cells (EMD:17034). **(F)** Superposition of the R27M NC spiral and helix maps. Scale bar, 50 Å in panels B–F. Panels B, C and F were made using the corresponding DeepEMhancer postprocessed maps at a contour level of 0.027, 0.07 and 0.07, respectively.

Importantly, our recent cryo-EM analysis demonstrated that insect cell-produced HRSV WT NC-like helices have a non-canonical symmetry, characterised by helical repetition of an asymmetric unit spanning ∼1.5 turns and composed of 16 N protomers, each adopting a distinct pose relative to the filament axis ^15^. Although the potential non-canonicity of the HMPV NC-like spirals was not explicitly investigated in the published work ^30^, re-examining the published medium-resolution cryo-EM map of ∼10 subunits per turn HMPV assemblies (Figure S4) reveals highly variable protomer tilt from the central axis. This feature hints at a possible non-canonical organisation of HMPV NCs, which may in turn be extrapolated to *Pneumoviridae* NCs in general. Surprisingly however, the cryo-EM maps of HRSV R27M spirals and filaments are visually more similar to regular *Paramyxoviridae* N-RNA assemblies ^39, 40, 41^. In addition (Figure 5, Figures S4, S5 and S6), the spirals and the filaments have the same structure at the level of both the individual protomer and the helical parameters, indicating that the spirals correspond to short or fragmented helices and are rigid enough to retain their pitch and twist in this short configuration.

The cryo-EM map of the R27M spirals has 2.53 Å average resolution and features a regular left-handed staircase of 9 protomers per turn instead of expected 10 (Figure 5B-C, Figure S5). The structures of individual R27M-N protomers and their relative arrangement remain quasi-identical upon local refinement of six or three central protomers (Figure S5). Moreover, each protomer has the same position and orientation relative to the spiral axis, demonstrating that, in contrast to the WT HRSV NCs and HMPV spirals, the HRSV R27M spiral reflects a canonical helical NC arrangement with a single R27M-N per asymmetric unit. The only notable variations between differently masked and locally refined maps focusing on smaller subsections of the spiral (Figure S5) are a slightly better definition of the outer rim of the NTD in the shorter subsections. These are likely attributable to minor variability in the relative tilting between the two subdomains (NTD and CTD) of R27M N (Figure 5B), as described for WT HMPV rings ^30^ and shown above by the MD simulations of an RNA-free HRSV N dimer that, in the case of R27M, has a tendency to remain in near-closed, RNA-bound-like states (Figure 4I).

As for the R27M NC-like helices (Figure 5D, Figure S6), in contrast to the WT NCs (Figure 5E), where the non-canonicity was already visible at the level of the 2D classes and their power spectra (PS) ^15^, the helical 2D classes of R27M NCs are clearly herringbone-shaped, with a ∼78 Å pitch and a near-integer number of subunits per turn (Figure S6A). Additionally, the PS exhibits a layer-line with the maximum close to the meridian at 1/78 Å, in agreement with the estimated pitch, while lacking a non-canonical layer-line indicative of a multi-protomer asymmetric unit (Figure S6A). This shows that the R27M-NC helix is composed of a single R27M-N protomer per asymmetric unit (Figure S4B) rather than 16 as in the WT NC. Interestingly, the ∼66 Å pitch calculated for a canonical helix closest to the non-canonical WT NC ^15^ matches the experimentally determined pitch of the C-terminally truncated N1-370 mutant (Figure 5F), which forms a canonical assembly due to the loss of periodic inter-turn interactions mediated by the CTD-arm ^15^. Remarkably however, the ∼78 Å pitch estimated from the 2D classes and PS of R27M-NC helices is ∼20% greater than that of WT NCs and identical to the pitch of the spiral assembly of R27M N (Figure 5G), confirming that the spirals are either formed by encapsidation of short RNA or by fragmentation of helical R27M NCs ^15^.

The cryo-EM map of the R27M NC helix (Figure 5D, Figure S6), calculated from ∼5 turn segments, has an average resolution of 3.1 Å, with 77.8 Å pitch and 8.97 subunits per turn, corresponding to a -40.14° twist and an 8.68 Å rise. These values should be compared to ∼10 subunits per turn and 6.58 Å rise of the simulated and experimental canonical helices from WT and N1-370 HRSV NCs produced in insect cells. Masking and local refinement of three consecutive protomers further improved the resolution to 2.4 Å (Figure S6B-C), again likely due to slight variations in the relative NTD-CTD tilting, as observed for HMPV N, and to the moderate long-range order of the helix. In comparison to the WT NCs, the spiral and helical assemblies of R27M NCs appear relatively rigid, which is particularly remarkable given that, with one fewer subunit per turn and a 36% greater rise, successive helical turns are much further apart, with no longitudinal contacts despite the presence of the full-length CTD-arm.

A closer look at the R27M NC structure readily explains this striking change in helical symmetry. The protomer of the R27M NC is tilted -62.5° relative to the filament axis (Figure S4), as the most lying protomers in the non-canonical WT NCs. Notably, while the RNA groove between NTD and CTD is unchanged between the WT and the R27M NC-like assemblies, both the NTD-arm harboring the R27M substitution and the CTD-arm undergo significant rearrangements (Figure 6A). In R27M NCs, the CTDs of the nucleoproteins, located on the inner face of the helix, draw closer to one another, while the visible parts of the C-terminal arms that lie on top of the CTDs of their downstream neighbours contract (Figure 6B-C, Supplementary movie 1). It remains unclear which of these changes drives the other — whether the CTDs converge and the C-terminal arms follow, or whether the C-terminal arms act like tweezers, pulling the CTDs together. In either case, this compaction of the interior correlates with a spread-out movement of the outer regions, particularly visible at the tips of the NTDs (Figure 6B-C, Supplementary movie 1).

**Figure 6:**
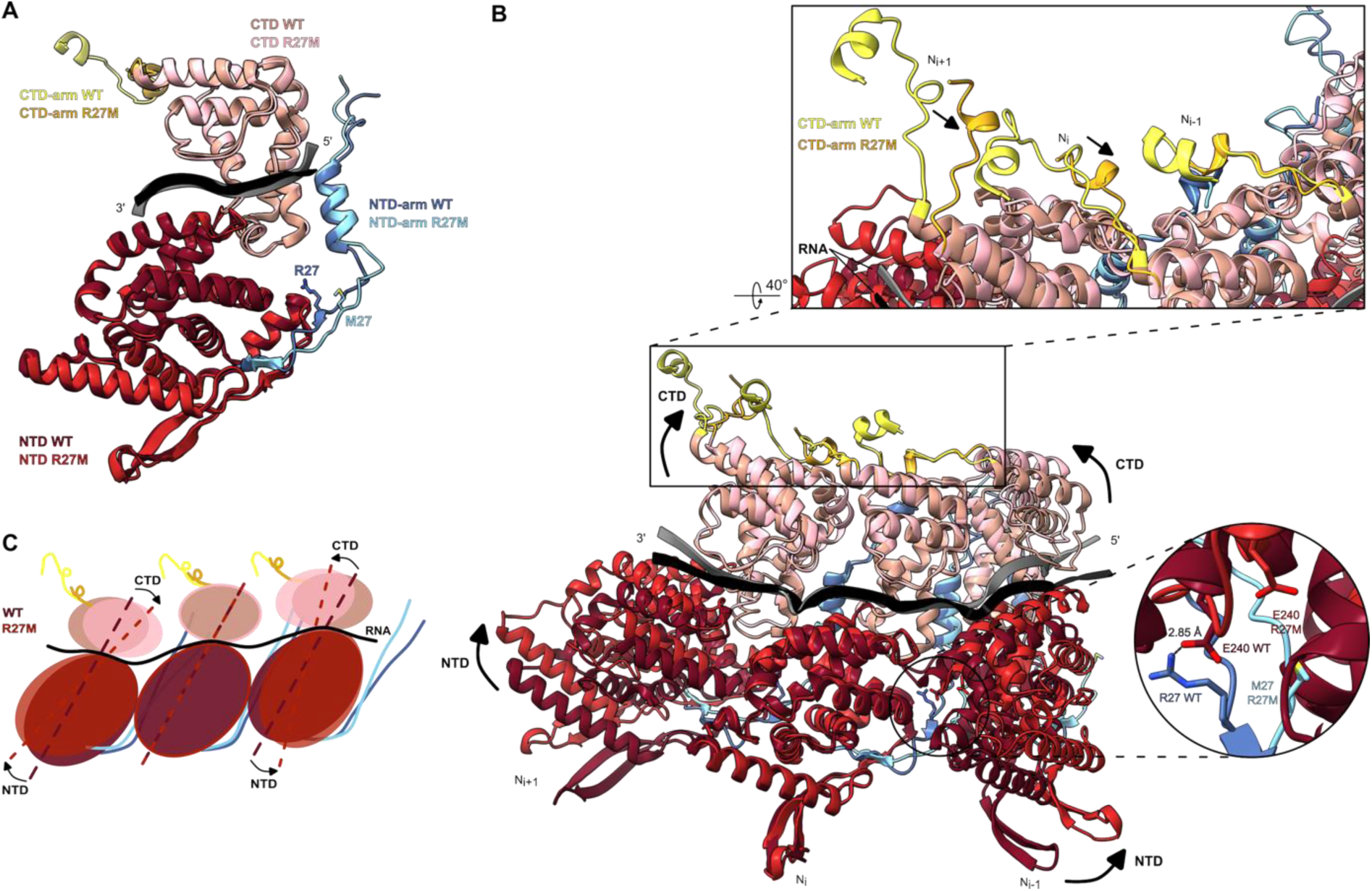
Lateral interactions in HRSV R27M NC. **(A)** Alignment of N protomers of the atomic models of the HRSV WT helical subsection (PDB: 8OP1) and the HRSV R27M NC. The NTD-arms, NTD, CTD and CTD-arms of HRSV WT helical subsection are coloured as in Figures 1 and 5. The RNA is in black. The HRSV R27M NC is coloured in light blue (NTD-arm), bright red (NTD), pink (CTD) and gold (CTD-arm). The RNA is in grey. **(B)** Alignment of 3 protomers from the atomic model of the HRSV WT helical subsection (PDB: 8OP1) and the HRSV R27M NC (the middle protomer is used for the alignment) revealing how in the R27M the CTDs are coming closer together and the NTDs pushed further apart in a fan-like movement compared to the WT NC (black arrows). Colouring same as in **A**. At the top, the close-up on CTD-arms is showing the contraction of the R27M NC CTD-arms compared to the ones from the WT NC. On the right side, the close-up is the R27-E240 interaction in the WT NC and its absence upon mutation of the residue 27 in the R27M NC. **(C)** Cartoon illustrating the fan-like movement of the protomers from the R27M NC compared to the WT NC. Colouring same as in **A**.

Unlike in the MD simulation of the RNA-free dimer, the R27-E240 interaction, which contributes to holding the NTDs together in the WT, completely disappears in the R27M NC. The overall motion resembles the opening of a fan: the center draws in as the periphery flares outward, or possibly the reverse (Figure 6B-C, Supplementary movie 1). This rearrangement is characterised by the disappearance of the WT HRSV-specific tripartite interaction between R234 of N_i-1_, D221 of N_i_ and Y23 of N_i+1_ in the so-called “N-hole” of the protomer N_i_, because Y23, located on the NTD-arm, close to the R27M mutation, shifts out (Figure S7). In sum, although the R27M mutation destabilises the R27-E240 and N-hole interactions, reduces the number of subunits per turn and increases the helical pitch of the R27M NC, all changes potentially expected to loosen the R27M helix in comparison to the WT, this methyl-mimetic mutation of R27 instead counterintuitively renders the R27M NC more rigid and straight. Taken together, our findings establish R27 as a key structural switch, regulating N-N interactions and conformational plasticity through post-translational modification, both in the RNA-free state and in the assembled N-RNA helix.

## DISCUSSION

In this study, we investigated the potential role of PTMs in regulating NC structural rearrangements and viral replication by conducting LC-MS/MS analyses of N, either immunoprecipitated from HRSV-infected human cells or purified from insect cells. The latter material had previously been used to solve the structure of the flexible helical WT HRSV NC, whereby we discovered its non-canonical symmetry ^15^. We chose to focus on PTMs found to overlap between both types of samples. Several PTMs were identified, clustered particularly on the NTD-arm of N, known to play a key role in N oligomerisation on the RNA ^15, 18, 42^. Interestingly, our recent study showed that phosphorylation of Y88 stabilises the NTD-arm of the monomeric, RNA-free, N^0^ form ^26^. The current findings therefore provide additional compelling evidence that the NTD-arm acts as a key regulatory hub controlling N activity via PTMs. Only three PTMs were detected in both analyses: phosphorylation of residues S20 and Y38, and dimethylation of R27 (Figure 1 and Table S1). Interestingly, none of these PTMs had been identified in our previous MS study of N rings produced in *E. coli* or of N^0^ purified from either *E. coli* or insect cells ^26^. This comparison, together with the previous observation that Y38 phosphorylation alters NC conformation ^29^, supports the idea that these PTMs are specific to assembled NCs. However, we cannot exclude that they may also depend on enzymatic activities specific to eukaryotic cells.

Our attention was specifically drawn to the dimethylation of R27 due to i) the high confidence score obtained from LC-MS/MS analysis, ii) its conservation among *Pneumoviridae* N proteins, and iii) its important role in the N-N interaction previously documented for HRSV and HMPV ^15, 30^. We demonstrated that R27A and R27M substitutions, which respectively mimic the removal or methylation of the arginine residue, impaired L activity and the formation of pseudo-VFs in transfected cells (Figure 2). Furthermore, the R27K substitution caused an approximately 90% reduction in polymerase activity without major impact on pseudo-VFs formation. The strong effects of these substitutions are consistent with the essential role of R27 in NC assembly, but also suggest that complete methylation of R27 is detrimental to L activity and VF morphogenesis. These results correlate with the low abundance of PTMs detected by LC-MS/MS, suggesting that either only a small fraction of protomers within each NC is methylated or that methylation occurs only on specific NC subsets, depending for instance on the accessibility of the NTD-arm to methyltransferases.

We next sought to determine whether R27 dimethylation corresponded to ADMA or SDMA ^33^. While immunoblot analyses using an anti-ADMA antibody failed to detect relevant labelling on purified N (Figure 3A and S1), the lack of commercial antibodies capable of detecting SDMA outside the RG/RGG motifs prevented direct assessment of SDMA. Nevertheless, the absence of ADMA detection, combined with the previously published observation that PRMT5, a type II arginine methyltransferase and main catalyst of SDMA, co-immunoprecipitates with N from transfected human cells ^35^, led us to hypothesise that R27 undergoes symmetric dimethylation mediated by PRMT5. Supporting this idea, our LC-MS/MS analysis identified a peptide with low-to-medium confidence score, carrying a mono-methylated R27. Indeed, PRMT5 is thought to function non-processively ^43^, allowing intermediate MMA products to accumulate. In contrast, PRMT1, the main driver of ADMA in eukaryotic cells, mediates dimethylation in a processive manner, whereby the mono-methylation step is immediately followed by the second methyl transfer ^33, 44^.

Having validated the interaction between PRMT5 and N in cells using a split-Nano luciferase assay, we further showed that silencing PRMT5 decreased overall SDMA levels in cell lysates and inhibited both HRSV replication and polymerase activity (Figure 3). Thus, our findings point to a delicate mechanism regulating NC function via R27 methylation. On one side, *in vivo* R27 dimethylation, most probably by PRMT5, enhances replication and polymerase activity; on the other, the R27M substitution induces a loss of both polymerase activity and of the ability to form LLPS with P. Altogether, our findings suggest that complete methylation rigidifies the NC and renders it non-functional, whereas partial methylation confers a functional advantage, likely through the notably increased flexibility of the non-canonical helical WT NC and the modulation of the RNA accessibility to the polymerase complex ^15^. The system appears finely tuned to precisely adjust N methylation in response to specific functional needs.

Although our data highlight the importance of PRMT5 in HRSV polymerase activity and replication, we were unable to detect PRMT5 within pseudo-VFs. Further investigation will thus be required to confirm its specific role in R27 methylation and/or to explore whether PRMT5 also mediates arginine methylation of other proteins of the HRSV replication complex, such as P, L or M2-1. Interestingly, our findings align with reports linking PRMTs to regulation of viral polymerases activity. Indeed, PRMT5 was shown to catalyse the methylation of the PB2 polymerase subunit of the avian influenza virus (IAV) ^36^ and of the VP1 polymerase of the infectious bursal disease virus (IBDV) ^45^, while PRMT1 was demonstrated to methylate the SARS-CoV-2 nucleoprotein ^46^.

Despite growing evidence for arginine methylation in prokaryotes ^47, 48, 49^, no PRMT5 analogue has been identified to date. To our surprise, while recombinant WT HRSV N-RNA purified from *E. coli* is widely known to form only rings, expression of the R27M mutant yields spirals and rigid, canonical helices. Considering the absence of R27 methylation in *E. coli*-derived rings ^26^, the R27M substitution appears to favour regular helical assembly. In this light, the non-canonicity of the HMPV N-RNA spirals produced in *E. coli* is intriguing and warrants further structural analysis of helical HMPV NCs expressed in either *E. coli* or insect cells, or purified from infected cells.

This brings us to the non-canonical NC organisation observed in insect cell-expressed HRSV N: could the variations in protomer tilt and shift within the asymmetric unit and the resulting increased NC flexibility represent structural fingerprints of distinct PTM states? Considering the corresponding variation in RNA accessibility along the non-canonical helix, it seems plausible that selective PTMs regulate RNA accessibility and thereby HRSV replication. The follow-up question is when exactly such modifications of the NTD-arm occur during the replication process? Taking into account i) the differences in PTMs identified on N^0^ versus assembled NCs, ii) the clustering of NC-specific PTMs at the NTD-arm of N, iii) the inaccessibility of this arm once buried inside the assembled NC, one can imagine a narrow window for modification, likely during or just after encapsidation, when a new N monomer binds the RNA emerging from the polymerase. At this stage, the NTD-arm of the newly incorporated subunit would already be engaging the preceding subunit but the connection is likely still loose, so that the arm could remain exposed to potential PTMs. Presumably, it is only when the next subunit arrives and locks the structure from the opposite side that this, now penultimate, NTD-arm becomes fully buried and inaccessible. This speculative mechanism is reminiscent of a zipper: as each tooth slots into place, it is not fully locked until the next one comes in and clamps it from the other side. The results of our MD simulation assessing the impact of dmR27 and the R27M mutation on an RNA-free N-N dimer are consistent with the idea of dimethylation occurring upon RNA encapsidation. Indeed, the simulations show that R27 dimethylation enhances opening and closing motions of the RNA binding groove of N while shifting the equilibrium towards the open state, which would favour RNA binding and encapsidation (Figures 4 and S3). Based on *in situ* observations that Ebola virus NCs evolve from loosely coiled to helical and eventually into bundles of rigid helices during infection ^50^, and given the rigidity of R27M NCs, one may hypothesise that newly synthesised HRSV NCs are more open than purified ones and thus accessible to local R27 dimethylation, which would trigger HRSV NC maturation. Both scenarios — dimethylation upon encapsidation or during maturation — may act in concert to fine-tune NC function ^50^.

Although MD simulations of an RNA-free nucleoprotein dimer cannot directly translate to the folding of an N-RNA helix, it suggests that the R-to-M substitution does not perfectly mimic symmetric R di-methylation: unlike dmR27, the R27M mutant tends to stay in a more closed conformation. Nevertheless, these simulations confirmed the crucial role of R27, and the R27M substitution remain the best available proxy to explore the structural and functional effects of R27 dimethylation. The cryo-EM structure of the R27M NCs revealed that modification of this NTD-arm residue induces long-range conformational changes, leading to significant rearrangements of the opposite, CTD-arm. We previously showed that the CTD-arm plays a critical role in NC organisation by periodically linking successive helical turns and conferring flexibility to the WT HRSV NC ^15^, and that its shortening also leads to rigid, canonical helices, albeit with distinct helical parameters (^15^ and Figure 5D-F). Taken together, our cryo-EM studies highlight that modulation of both the NTD- and CTD-arms profoundly impacts the helical NC organisation and flexibility, revealing the strong structural interplay between these two terminal domains of N. It is tempting to speculate that NC conformational changes modulating RNA accessibility may also depend on other PTMs targeting the NTD- and CTD-arms. Further investigation of these PTMs, their potential combinations and their temporal regulation may shed light not only on the mechanism of HRSV genome replication, but also on the NC rearrangements underlying the transition from replication to NC transport to the cell plasma membrane and incorporation into new virions.

## Acknowledgements

We thank Mrs Pierre Adenot, Matthieu Simion and Vlad Costache for access to the MIMA2 platform (Microscopie et Imagerie des Microorganismes, Animaux et Aliments, https://doi.org/10.15454/1.5572348210007727E12), and their assistance with the confocal microscopy. Molecular graphics images were produced using Chimera and PyMol softwares. This work was carried out with the financial support of the French Agence Nationale de la Recherche, specific program ANR RSVFact (ANR-21-CE15-0030-02), ANR SofteN (ANR-23-CE11-0027-01) and ANR PneumoPEPS (ANR-23-CE11-0027-01). We thank Guy Schoehn for establishing and managing the IBS cryo-EM platform and for providing training and support, and Lefteris Zarkadas for assistance at the Glacios microscope. For cryo-EM grid preparation and screening, we used the platforms of the Grenoble Instruct-ERIC centre (ISBG; UAR 3518 CNRS-CEA-UGA-EMBL) within the Grenoble Partnership for Structural Biology (PSB), supported by FRISBI (ANR-10-INBS-0005-02) and GRAL, financed within the University Grenoble Alpes graduate school (Ecoles Universitaires de Recherche) CBH-EUR-GS (ANR-17-EURE-0003). The IBS EM facility is supported by the Rhône-Alpes Region, the Fondation pour la Recherche Médicale (FRM), the fonds FEDER and the GIS-Infrastructures en Biologie Sante et Agronomie (IBISA). We thank the FRISBI access scheme (9^th^ access call), Nils Marechal and Alexandre Durand for enabling collection of the final cryo-EM dataset at a Krios G4 microscope of the Integrated Structural Biology platform of the Strasbourg Instruct-ERIC center IGBMC-CBI supported by FRISBI (ANR-10-INBS-0005) and EquipEx+ France cryo-EM (ANR-21-ESRE-0046). We acknowledge Umeå Centre for Electron Microscopy (UCEM) for technical assistance. Support was provided by SciLifeLab Integrated Microscopy Technology Unit at Umeå University and the Swedish Research Council (VR) Tage Erlander visiting professorship Grant (DNR 2022-00308) to IG.

## Author contributions

V.B., J.-F.E., I.G. and M.G. designed experiments. V.B. and M.G performed cellular and biochemical assays. J.S. performed microscopy analysis. L.G., C.S.S., M.B.-V., A.D. and I.G. performed cryo-EM analysis. C.S.S. built atomic models. C.L. performed molecular dynamics simulations. C.R.-M. and C.H. performed mass spectrometry analysis. I.G. and M.G. wrote the paper with input from V.B. and C.L. L.G., C.S.S., I.G. and M.G. edited the manuscript. All authors commented on the manuscript.

## Material and methods

### Cells

BHK-21 cells (clone BSRT7/5) constitutively expressing the T7 RNA polymerase ^51^, A549 and HEK293T cells were grown in Dulbecco’s modified Eagle’s medium (Eurobio Scientific, Les Ulis, France) supplemented with 10% fetal calf serum (FCS), 2 mM L-Glutamine, and antibiotics. HEp-2 cells were grown in Eagle’s MEM (Eurobio Scientific, Les Ulis, France) supplemented with 10% FCS, 2 mM L-Glutamine and antibiotics. Cells were transfected using Lipofectamine 2000 (Invitrogen), and/or Lipofectamine RNAiMAX (Invitrogen) as described by the manufacturer.

### Plasmids & bacmids

As previously described, the codon-optimized sequence coding for the wild type (WT) N (strain Long), cloned in the pFastBacDual vector under the control of the polyhedrin promoter, between BamHI and SalI sites (pFBD-N) ^15^ was used for N expression in insect cells. Briefly, for bacmid generation, DH10EmBacY cells were transformed with pFBD-N. After overnight incubation at 37°C, plates were stored at 4°C for 2-3 days, and the colonies that remained white were selected for bacmid obtention, using the NucleoBond BAC100 kit (Macherey-Nagel). Plasmids for eukaryotic expression of the HRSV N, P, M2-1 and L proteins, designated pN, pP, pM2-1 and pL, and the pM/Luc subgenomic minigenome which encodes the firefly luciferase (Luc) reporter gene under the control of the M/SH gene start sequence have been described previously ^31, 52^. Commercially available pciNanoLuc 114 and 11S vectors (GeneCust) were used to clone the HRSV A2 codon-optimized P, N, and PRMT5 constructs using standard PCR, digestion and ligation techniques. For production of recombinant proteins in *E. coli*, the previously described pET-N and pGEX-PCT plasmids were used ^20, 22^. Mutations were generated using the Q5 Site-Directed Mutagenesis Kit (New England Biolabs), following the manufacturer’s instructions. Sequence analysis was carried out to check the integrity of all the constructs.

### Antibodies

The following primary antibodies were used for immunofluorescence and/or immunoblotting: a rabbit anti-N antiserum ^53^, mouse monoclonal anti-N antibody (Serotec), a rabbit anti-P antiserum ^28^, a mouse monoclonal anti-P ^54^, a mouse monoclonal anti-PRMT5 (Santa Cruz Biotechnology), a mix of rabbit monoclonal anti-adme-R (Cell Signaling Technology), a mix of rabbit monoclonal anti-sdme-RG (Cell Signaling Technology), and a mouse monoclonal anti-β-tubulin (Sigma). Secondary antibodies directed against mouse and rabbit IgG coupled to HRP (SeraCare) were used for immunoblotting, and antibodies directed against mouse and rabbit IgG coupled to Alexa 568 or Alexa 488 (Invitrogen) were used for immunostaining.

### Production and purification of recombinant proteins

For production of N recombinant protein in *E. coli*, BL21 (DE3) bacteria were co-transformed with the pET-N and pGEX-PCT (for expression of GST fused with the C-terminal residues of P, residues 161-241) plasmids, as previously described ^28^. Cultures were grown at 37°C in 2xYT medium containing 100 µg/mL ampicillin and 50 µg/mL kanamycin. After 8h, the same volume of 2xYT was added to the cultures, and protein expression was induced by adding 80 µg/mL IPTG overnight at 28°C. Bacteria were harvested by centrifugation, and pellets resuspended in 15 mL of TEN lysis buffer (50 mM Tris-HCl, 10 mM EDTA, 150 mM NaCl, pH 7.4, 0.6% NP-40 (v/v), anti-proteases, anti-phosphatases (Roche), RNase A (200 µg/mL, Invitrogen), DNase I (Promega). Resuspended pellets were incubated at 37°c for 40min, and supernatants were further collected after 12,000g centrifugation for 20min at 4°C, then incubated for 3h at 4°C with GST-PCT beads that were previously washed three times in lysis buffer. Thrombin cleavage was conducted overnight at 4°C, and supernatants containing the purified recombinant N proteins were collected (Figure S3). The presence of RNA associated to WT and R27M N proteins was further confirmed by the OD_260_/OD_280_ ratio of 1.5, and by analysis of purified proteins on a native agarose gel and UV exposure, as well as native acrylamide gel analysis (Figure S3). For eukaryotic production of NC, High-Five cells (ThermoFisher Scientific, catalogue number B85502) were infected at MOI 2 with the baculovirus encoding an optimized sequence of N, for 72h ^15^. Cells were centrifuged at 3000g for 5 min and resuspended in 15 mL of TEN lysis buffer. Purification was then performed following the same protocol as described above, by affinity for GST-PCT beads.

### Purification of N from infected cells

HEp-2 cells infected for 24h with rHRSV-mCherry at MOI 3 were lysed for 30 min at 4°C in ice-cold lysis buffer (50 mM Tris-HCl pH 7.4, 2 mM EDTA, 150 mM NaCl, 0.5% NP-40) with a complete protease inhibitor cocktail (Roche) and phosphatase inhibitor cocktail (Roche), then centrifuged for 10 min at 12,000g. Supernatant was incubated for 4h at 4°C with a mouse monoclonal anti-N antibody coupled to agarose beads (Euromedex). The beads were then washed 3 times with lysis buffer and 1 time with PBS, and proteins were eluted in Laemmli buffer at 95°C for 5 min and then subjected to SDS-PAGE.

### Mass spectrometry analysis

Samples were loaded on an SDS-PAGE gel (NuPAGE 4-12% Bis-Tris gel, Invitrogen), and each strip of interest was cut and washed successively for 15 min with a solution of (i) 10% acetic acid and 40% ethanol (ii) acetonitrile (ACN) and 50 mM ammonium bicarbonate mixture (1:1). Then, disulfide bonds were reduced with dithiothreitol 10 mM at room temperature during 30 min and alkyled with iodoacetamide 55 mM during 45 min in the dark at room temperature. Digestion was performed in 50 mM ammonium bicarbonate (pH 8.0) with a protein:enzyme ratio of 1:100 (trypsin, AspN and chymotrypsin), overnight at 25°C for trypsin, AspN, and chymotrypsin. Peptides were extracted by adding 0.5% TFA / 50% ACN solution, the supernatant was lyophilized and stored at -20°C. Lyophilized peptides were then resuspended in 20-40 µL of 2% ACN and 0,08% TFA buffer and MS analyses were performed on a Dionex U3000 RSLC coupled to an Orbitrap Fusion™ Lumos™ Tribrid™ mass spectrometer (Thermo Fisher Scientific) at the Plateforme d’Analyse Protéomique de Paris Sud Ouest (PAPPSO), Jouy-en-Josas, France (http://pappso.inrae.fr/).

The tryptic peptides were loaded on a PepMap Neo trap column (300µm i.d. x 5mm, with a particle size of 5 µm, 100A, Thermo Fisher) and separated using a C18 column (50 cm x 75 µm i.d.2 µm particle size). The peptides were eluted on the nLC system through the following gradient elution program: 2.5-35 % buffer B (80% ACN, 0.1% FA) within 0-50min, 35-45 % buffer B in 50-55min, 45-98% buffer B in 55-57 min. The detection of peptides was acquired in the DDA mode, for MS1 signals, the electrospray voltage was set at 1600V, in positive mode. MS scans were performed at 120,000 resolution, m/z range 400 ‒ 2000 Da. LC-MS/MS analysis was performed in a data-dependent acquisition (DDA), with a top speed cycle of 2.5 s for the most intense double or multiple charged precursor ions. Ions in each MS scan over threshold 20,000 were selected for fragmentation (MS2) by higher energy collisional dissociation (HCD) for N purified from infected HEp-2 cells and electron transfer higher energy collisional dissociation (ETHCD) for recombinant NC purified from Hi5 cells, at 30% for identification and detection in the orbitrap followed by a top speed MS2 fragment ion. Precursors were isolated in the quadrupole with a 1.6 m/z window and dynamic exclusion within 10 ppm during 60 s was used for m/z-values already selected for fragmentation. The AGC targets are fixed as standard for MS and MS/MS analysis. Polysiloxane ions m/z 445.12002, 519.13882 and 593.15761 were used for internal calibration.

### Protein identification from LC-MS/MS

Protein identification was performed using Peaks Studio 10.6 software. A FASTA format database was used including the genomes of uniprotkb_Trichoplusia_Ni (20889 entries) or *Homo Sapiens* (20403 entries), the N_HRSV protein and the common contaminants proteins. We used PTMs parameters, focusing on phosphorylation and methylation modifications but oxidation of methionine and acetylation were also selected as potential modifications. Filters parameters were set at 10 ppm for parent mass error tolerance, 0.02 Da for fragment mass error tolerance and a maximum of 3 missed cleavages were fixed.

### Minigenome assay

Cells at 90% confluence in 96-well dishes were transfected with a plasmid mixture containing 62.5 ng of pM/Luc, 62.5 ng of pN, 62.5 ng of pP, 31.25 ng of pL, and 15.5 ng of pM2-1 as well as 15.5 ng of pRSV-β-gal (Promega) to normalize transfection efficiencies ^31^. Transfections were done in 5 replicates, and each independent experiment was performed three times. Cells were lysed 24 hours after transfection in luciferase lysis buffer (30 mM Tris, pH 7.9, 10 mM MgCl2, 1 mM DTT, 1% Triton X-100, and 15% glycerol). The luciferase activities were determined for each cell lysate with an Infinite 200 Pro (Tecan Infinite M200PRO, Männedorf, Switzerland) and normalized based on β-galactosidase (β-gal) expression. N and tubulin expression was analyzed by immunoblotting in each experiment.

### NanoLuc interaction assay

Constructs using the NanoLuc subunits 114 and 11S were used ^37^. HEK293T cells were seeded in 48-well plates, at a concentration of 3.10^4^ cells per well. After 24h, cells were co-transfected with 0.4 µg of total DNA (0.2 µg of each plasmid), using Lipofectamine 2000 (Invitrogen). 24h post-transfection, cells were washed once with PBS 1X and further lysed for 1h at room temperature in 50 µL of NanoLuc lysis buffer (Promega, Madison, WI, USA). The enzymatic activity of the Nano Luciferase was measured using the NanoLuc substrate (Promega). For each pair of plasmids, Normalized Luminescence Ratios (NLR) were calculated as follows: the luminescence activity measured in cells transfected with the two plasmids (each viral protein fused to a different NanoLuc subunit) was divided by the sum of the luminescence activities measured in both control samples (each NanoLuc fused viral protein transfected with a plasmid expressing only the NanoLuc subunit). Luminescence was measured using the Infinite 200 Pro (Tecan Infinite M200PRO, Männedorf, Switzerland). For each condition, 6 technical replicates were performed, and the experiment was repeated 4 times.

### Fluorescence microscopy

Cells grown on coverslips were transfected with pN (WT or mutant) and pP. Twenty-four hours after transfection, cells were fixed with 4% paraformaldehyde (PFA) for 20 min. Fixed cells were permeabilized, blocked for 30 min with phosphate-buffered saline (PBS) containing 0.1% Triton X-100 and 3% bovine serum albumin (BSA), and then successively incubated for 1h at room temperature with primary and secondary antibody mixtures diluted in PBS containing 3% BSA. For labeling nuclei, Hoechst 33342 (Invitrogen) was added during incubation with the secondary antibodies. Coverslips were mounted in Prolong gold antifade reagent (Invitrogen). Cells were observed with a Zeiss LSM 700 confocal laser scanning microscope (MIMA2 Platform, INRAE), and images were processed using Zen 2011 and ImageJ software.

### SDS-PAGE and Western Blots

Protein samples were separated by electrophoresis on 12% polyacrylamide gels in Tris-glycine buffer. All samples were boiled for 3 min prior to electrophoresis. For Western blot, proteins were then transferred to a nitrocellulose membrane (BioRad). The blots were blocked with 5% nonfat milk in PBS Tween20 0.2%, followed by incubation with primary, then secondary antibodies. Western blots were developed using ClarityTM Western ECL substrate (Bio-Rad) and exposed using the Bio-Rad ChemiDoc Touch Imaging System (Bio-Rad, Hercules, CA, USA).

### Virus infection

The recombinant HRSV Long strain expressing the mCherry (rHRSV-mCherry) was used for infection experiments ^27^. For infection of HEp-2 and A549, cells were incubated in the presence of virus at a multiplicity of infection (MOI) of 1 and 0.2 respectively, for 2 h in MEM or DMEM without phenol red and without FCS. The medium was then replaced with medium containing 2% FCS, and plates were incubated for 24h or 48h at 37°C with 5% CO_2_. The mCherry fluorescence was measured using a spectrophotometer (Tecan Infinite M200PRO, Männedorf, Switzerland) with excitation and emission wavelengths of 580 and 620 nm, respectively.

### siRNA experiments

The control siRNA (siCT) and the siRNA against PRMT5 (siPRMT5) were purchased from ThermoFisher Scientific (Waltham, MA, USA). The PRMT5 siRNA is a pool containing 6 different sequences (RefSeq NM_001039619.2, NM_001282953.1, NM_001282954.1, NM_001282955.1, NM_001282956.1, NM_006109.4). A549 or BSRT7/5 cells were transfected with the indicated siRNAs at a final concentration of 10 nM by reverse transfection into 96-well plates (7.10^4^ cells per well) using Lipofectamine RNAiMAX (ThermoFisher Scientific), according to the manufacturer’s instructions. Briefly, a mixture containing Opti-MEM (Invitrogen), Lipofectamine RNAiMAX and the desired siRNA was incubated at room temperature for 5 min before being deposited at the bottom of the wells. A549 or BSRT7/5 cells were then added drop by drop before incubation at 37°C with 5% CO_2_. Potential cytotoxicity of siRNA treatment was assessed using the CellTiter-Glo luminescence cell viability kit (Promega, Madison, WI, USA).

### Statistical analysis

A nonparametric Mann-Whitney test (comparison of two groups) was used to compare unpaired values (GraphPad Prism software). Significance is represented in the figures (*, *P* < 0.05; **, P < 0.01)

### Molecular Dynamics Simulations

Classical MD simulations were used to study the impact of Arg27 modification (dimethylation or mutation to methionine) on the dynamics of N-N interactions. Dimeric N models were extracted from a 2.8 Å resolution cryo-EM structure of HRSV N-RNA rings (PDB code 80P2) ^15^ or from the R27M structure. MD systems corresponding to WT R27, dmR27 or R27M dimers were set up using the solution builder tool from CHARMM-GUI input generator ^55^. Briefly, each system was solvated in a rectangular periodic box, which size was determined by the biomolecular extent, resulting in the addition of approximately 65k TIP3P water molecules. The systems were neutralized by adding 150mM NaCl. Each three systems were then energy minimized, equilibrated in NVT ensemble and simulated for at least 700ns in 3 independent trajectories in GROMACS2024 ^56^ using CHARMM-GUI scripts ^57^. The CHARMM36m force field was used for all simulations ^58^. The trajectories were analysed using GROMACS tools to extract snapshots, inter-residue distances, and to perform a PCA analysis on the CA atoms of the N_i_ protomers stripped of the NTD and CTD arms, in order to analyse global conformational changes induced by R27 modifications.

### Cryo-EM grid preparation, data collection and preprocessing

For cryo-EM grid preparation, a 3.5 µl drop of purified R27M N-RNA sample was applied to a glow-discharged R2/1 300 mesh holey carbon copper grid (Quatifoil Micro Tools GmbH) and plunge-frozen in liquid ethane using a Vitrobot Mark IV (Thermo Fisher Scientific) operated at room temperature at 100% humidity. Grids were pre-screened on a Glacios microscope (Thermo Fisher Scientic) of the EM platform of the IBS, Grenoble, France. The final dataset was collected at the EM platform of the IGBMC, Strasbourg, France, on a Titan Krios G4 microscope (Thermo Fisher Scientic) with C-FEG, operating at 300 kV with a Selectris X energy filter (slit width 10 eV). Data was recorded on a Falcon 4i direct electron detector (Thermo Fisher Scientic), using the EPU software (Thermo Fisher Scientific), at a magnifcation of ×165,000 corresponding to a pixel size of 0.729 Å/pixel at the specimen level, with a defocus range of -0.8 to -2.2 µm in steps of 0.2 µm, a dose rate of 13.77 41 e-/Å^2^/s and a 3 second exposure, corresponding to a total dose of 41 e-/Å^2^. 9 images were acquired per hole. See Table S2 for a summary of data collection information.

A total of 20,988 movies were acquired, motion-corrected, dose-weighted, and CTF estimation performed on the aligned and dose-weighted summed frames using cryoSPARC 4.7.0 ^59, 60^. After manual inspection of the resulting micrographs, 20,495 were selected, discarding those with significant ice contamination and visible beam-induced movement. Further inspection of the micrographs showed the presence of short spiral filaments as well as herringbone-shaped helical filaments. Interestingly, although cryo-EM images of *E. coli* produced HMPV N-RNA ^30^ contained mostly flat rings (with a C10, sometimes C11, and rarer C12 symmetry), we did not observe flat 10, 11 or 12-mer views. However, we noticed occasional seemingly C9-symmetrical top views possibly corresponding to flat 9-subunit rings. We did not attempt to reconstruct these rings due to the lack of clear side views, but first and foremost because we chose to prioritise the more biologically relevant spiral and helical assemblies. The following image analysis was carried out in cryoSPARC 4.7.0 ^59, 60^.

### Cryo-EM image analysis of the R27M-NC spirals

An initial manual picking of 533 spirals, on a subset of micrographs, was performed to generate 2D classes. The selected 2D classes were used as template to perform automatic picking on all the micrographs: 1,310,353 particles were picked. The particles were selected by 2D classification iteratively until reaching a subset of particles used for *ab initio* reconstruction and then non-uniform refinement giving an initial map (212,400 particles) with a resolution of approximately 3.5 Å. The map was used to create projections for two sets of templates and do two automatic pickings in parallel: “top views” and “side views” pickings. This resulted into 1,106,242 and 1,050,155 particles for the top and side views pickings, respectively. Extensive 2D classification and selection resulted into subset of 229,847 and 649,908 particles of top and side views of the spirals, respectively. After removing the duplicate particles (2,185), the subsets of particles were merged followed by 3D classification using the volume of previous non-uniform refinement in order to remove the helices hidden in the side views of the spirals. A final non-uniform refinement resulted into 3D reconstruction (312,385 particles) displaying an average resolution of 2.53 Å (Fourier Shell Correlation (FSC) at 0.143), sharpened with a B-factor of -69.7 Å^2^. Masks enclosing three and six N protomers were created from the spiral map using RELION 4.0.1 ^61^ and imported into cryoSPARC to perform local refinements. The two final maps had an average resolution of 2.59 Å (six-protomers subsection) and 2.48 Å (three-protomers subsection) (FSC at 0.143) and were sharpened with a B-factor of -63.4 and -65.1 Å^2^, respectively.

### Cryo-EM image analysis of the R27M-NC helices

For the R27M-NC helices, an initial manual picking of 473 filaments was done on a subset of micrographs and used to generate 2D classes, subsequently used for automatic picking with the cryoSPARC filament tracer. Box size of 320 pixels and 24 pixels overlap between successive boxes were used, leading to extraction of 1,967,024 particles, which were subjected to 2D classification. While some of the classes clearly corresponded to double-headed or ring-capped helices, as previously observed for the WT NCs ^15^, we chose not to proceed with their 3D analysis but to focus on the most important, continuous helices only. After extensive 2D classification, 938,469 particles were selected. The average power spectra (PS) of several 2D classes were calculated (Figure S6) and inspected using e2display.py ^62^. All averaged PS displayed a layer line with a maximum close to the meridian at ∼1/78 Å corresponding to the estimated ∼78 Å helical pitch visible on the 2D classes. The helical twist was determined by running in parallel several helical refinements with a fixed pitch of 78 Å and a range of N protomers varying from 9 to 11 per turn. The best map displayed a helical pitch of 77.824 Å with 8.969 N protomers per turn (twist = -40.141° and rise = 8.677 Å) and an average resolution of 3.14 Å (FSC at 0.143) and sharpened with a B-factor of -75.9 Å^2^. The spiral map is almost perfectly fitting into the helical map. A mask enclosing three N protomers was created from the helical map using RELION and imported into cryoSPARC to perform local refinement. The three protomers subsection map had an average resolution of 2.39 Å (FSC at 0.143). The map was sharpened with a B-factor of -64.7 Å^2^.

### Atomic model building

The final B-factor-sharpened map obtained for the three-protomer subsection of the R27M helix was sharpened or blurred in Coot 0.9.8.95 ^63^ and used for model building. For this, three copies of the N-RNA protomer coming from the non-canonical WT NC five-protomer subsection map (PDB: 8OP1, chain A ^15^) were used as template and rigid-body-fitted into the map using ChimeraX (1.9rc202411061937) ^64^. The arginine 27 residue (R27) was changed into a methionine (M27) and the generated model was then refined into the map in an interactive manner with Ramachandran, secondary structure, base-pair and base-stacking restraints, using both Coot and Phenix software packages (1.21.2-5419) ^65^. The middle protomer (N_i_ protomer, termed Chain A), was then used for fitting into the equivalent protomer of the B-factor-sharpened cryoSPARC map obtained for the R27M NC helix (monomer 11, also named chain A), and a total of 25 monomers were generated in ChimeraX, applying the corresponding helical parameters. Adequate fitting into the helical map was confirmed using both Chimera and Coot.

## Supplementary Table and Figure legends

**Table S1:**
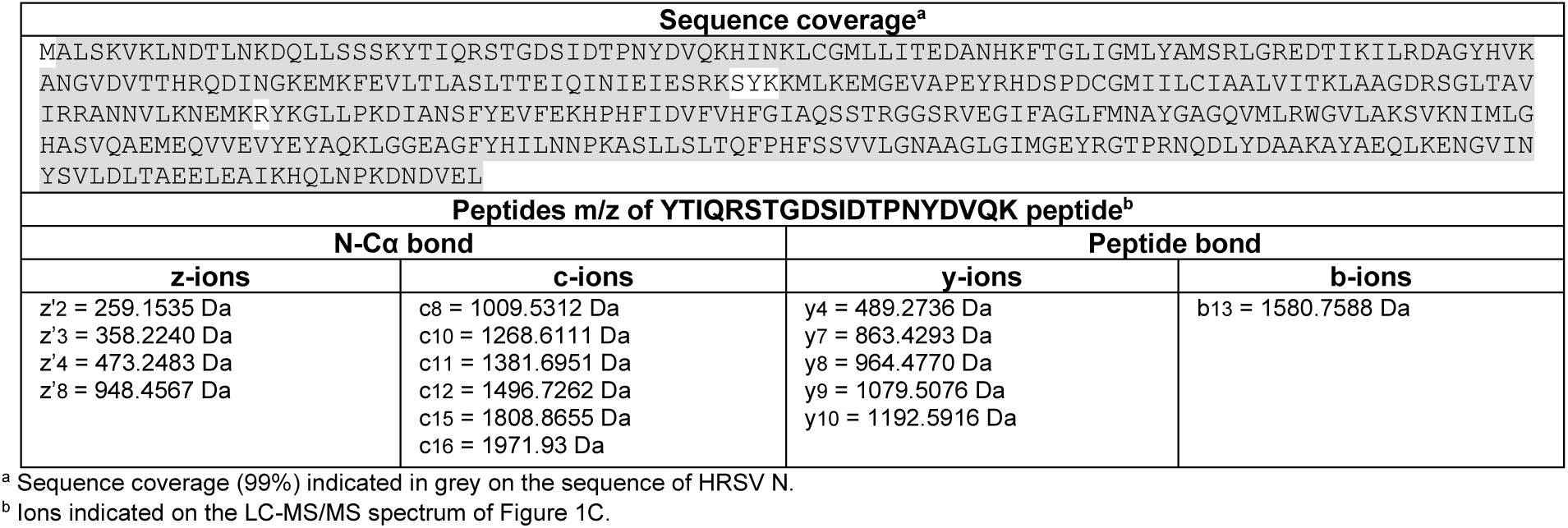
LC/MS-MS analysis of the N protein purified from infected cells, and of the di-methylated peptide.

**Table S2:**
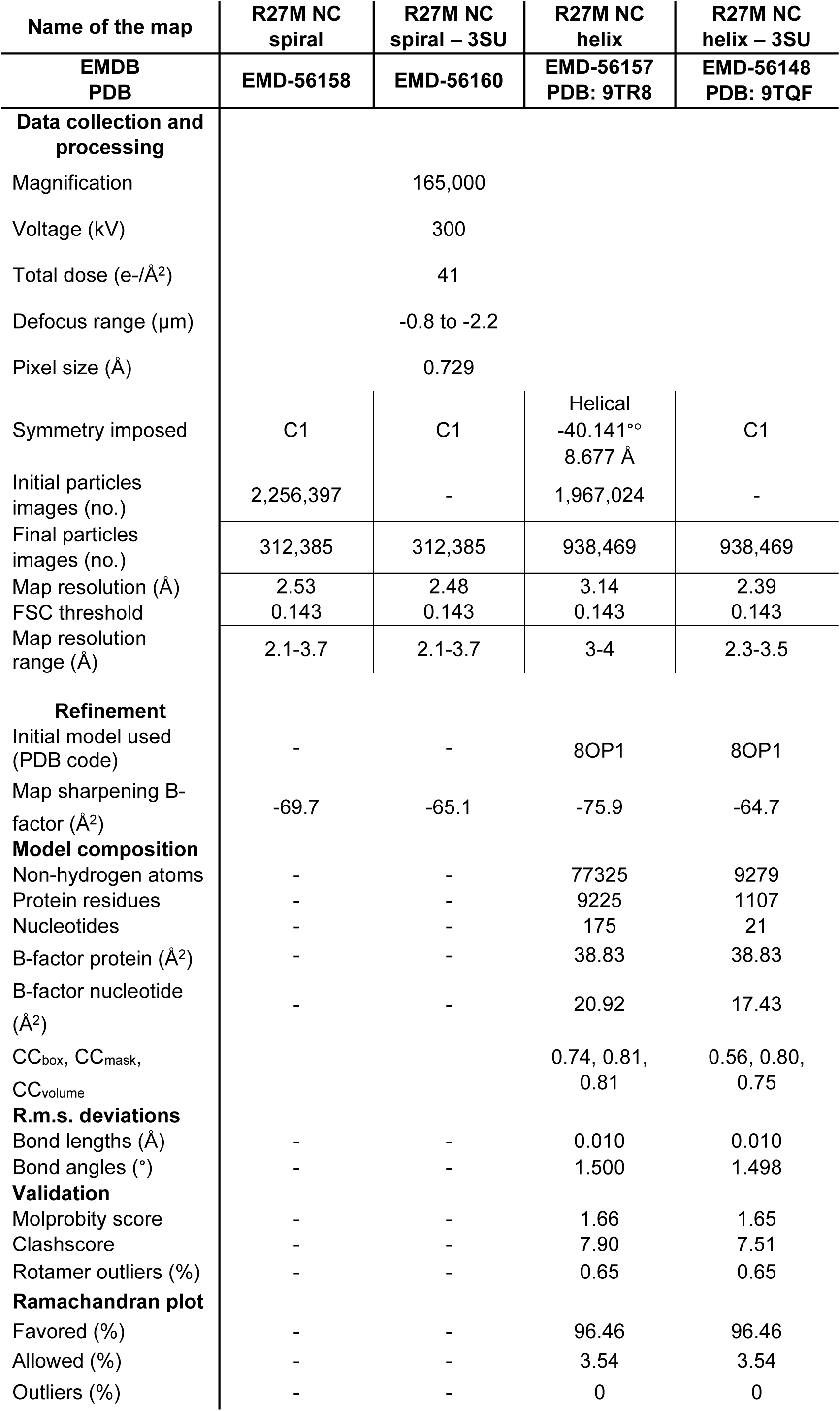
Cryo-EM data collection, refinement and validation statistics.

**Figure S1:**
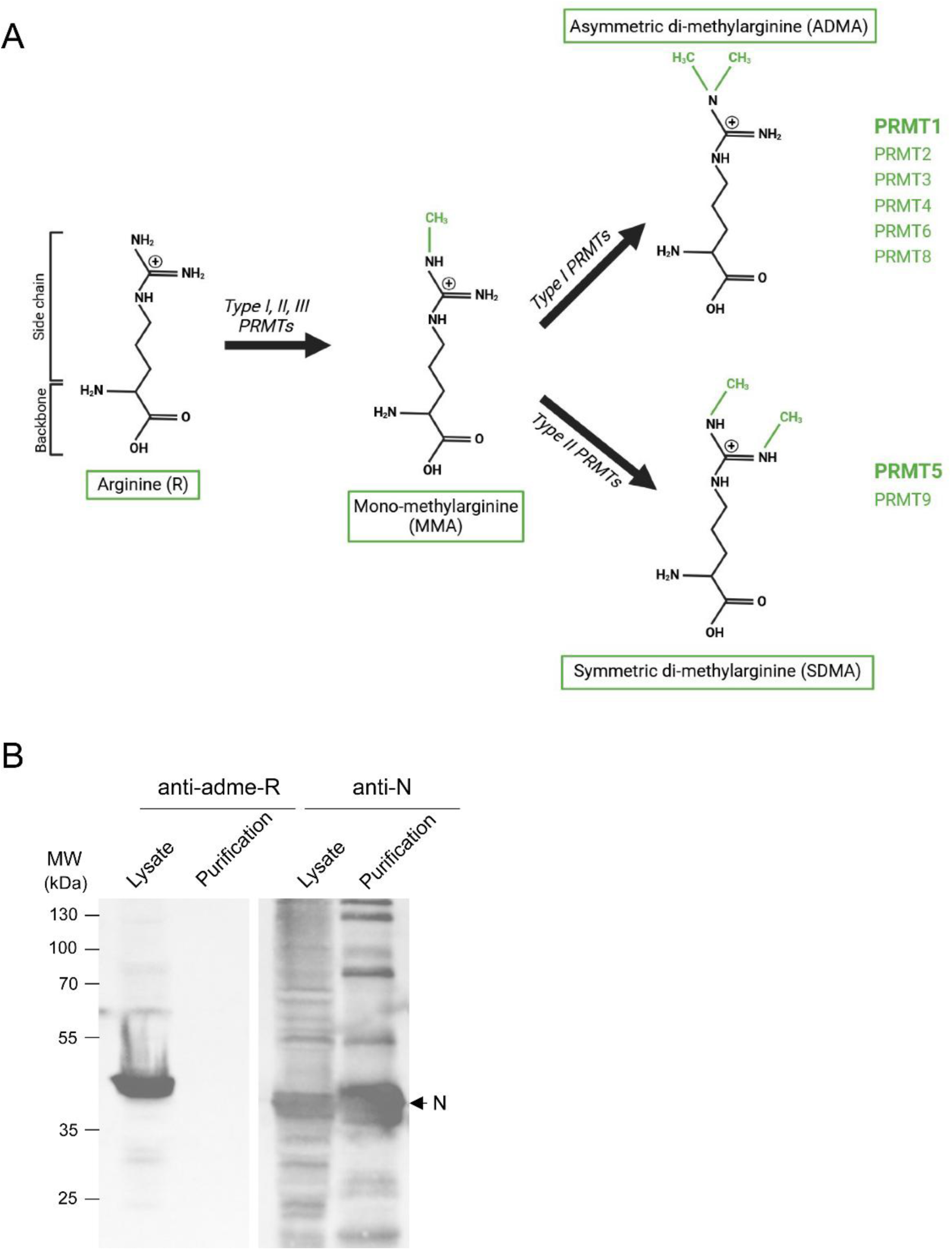
Study of the nature of the arginine dimethylation of N. **(A)** Schematic of the process of arginine dimethylation and of the PRMTs responsible of asymmetric and symmetric dimethylation. Added methyl groups are indicated in green. PRMT1 and 5, the most abundant PRMTs of each, type I and II PRMTs respectively, are indicated in bold. Adapted from ^66^ with BioRender.com. **(B)** Western blot analysis of the profile of proteins with ADMA in the lysate of High-Five cells expressing the HRSV N protein, and on corresponding purified recombinant N.

**Figure S2:**
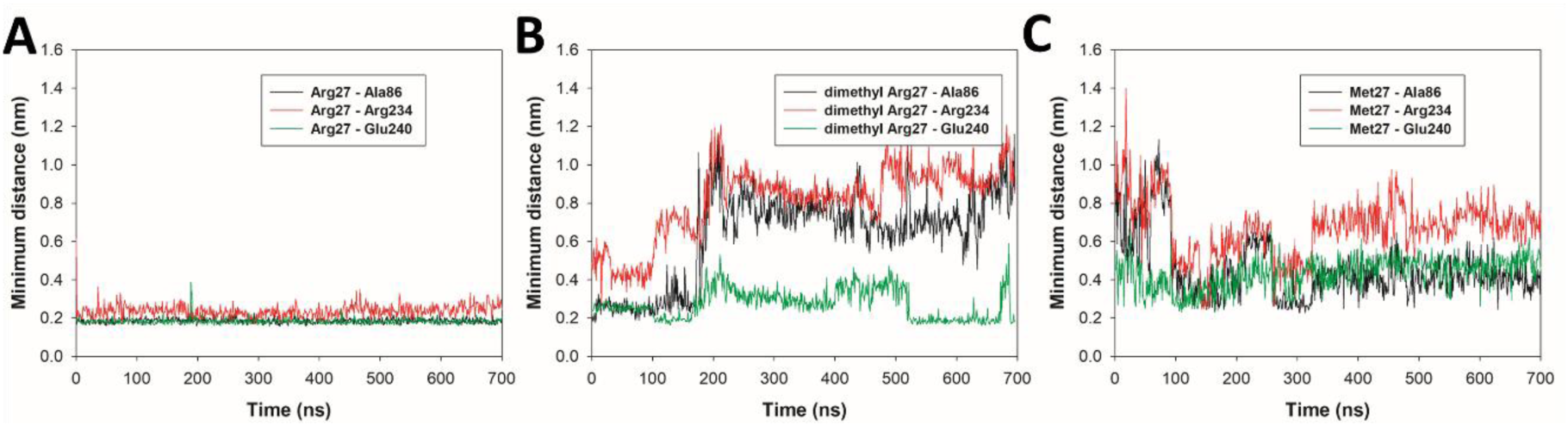
Minimum distances as a function of simulation time between R27 and interacting residues. A86, R234 and E240 in unmodified R27 **(A)**, dmR27 **(B)** and R27M **(C)** systems, highlighting the disruption of key canonical interactions in dmR27 and R27M.

**Figure S3:**
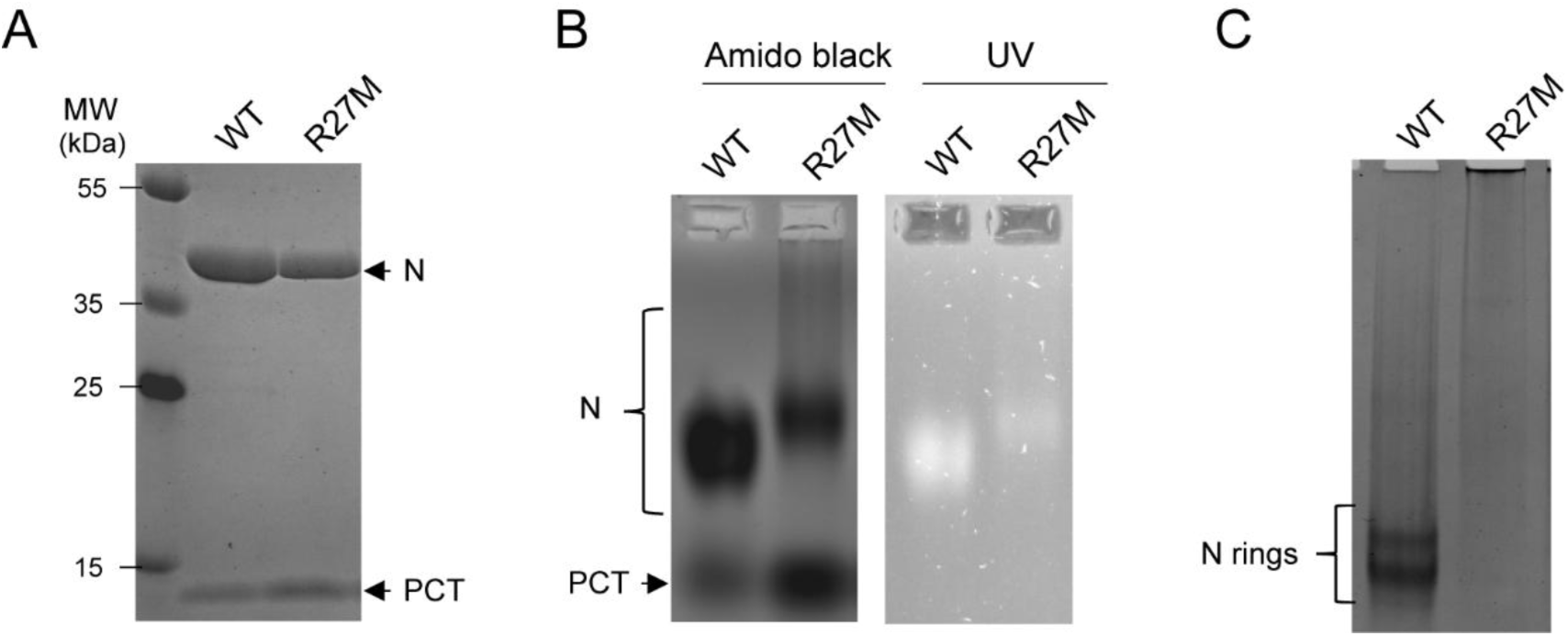
Analysis of purified R27M mutant of N. **(A)** Recombinant WT and R27M N proteins were expressed in *E. coli* and purified by affinity with GST-PCT. After cleavage of the GST by incubation in the presence of thrombin, the eluted samples were analysed by SDS-PAGE coloured with Coomassie blue. **(B)** Analysis of the purified proteins by migration on native agarose gel, showing a difference in migration profiles between WT and R27M proteins. Proteins were revealed by coloration with amido black, and the presence of RNA was observed by exposition of the gel to UV. **(C)** Analysis of the samples by migration on native acrylamide gel which reveals a smear for the R27M mutant, compared to the WT N that displayed the characteristic two-band pattern corresponding to the previously described 10-N and 11-N oligomers ^18^.

**Figure S4:**
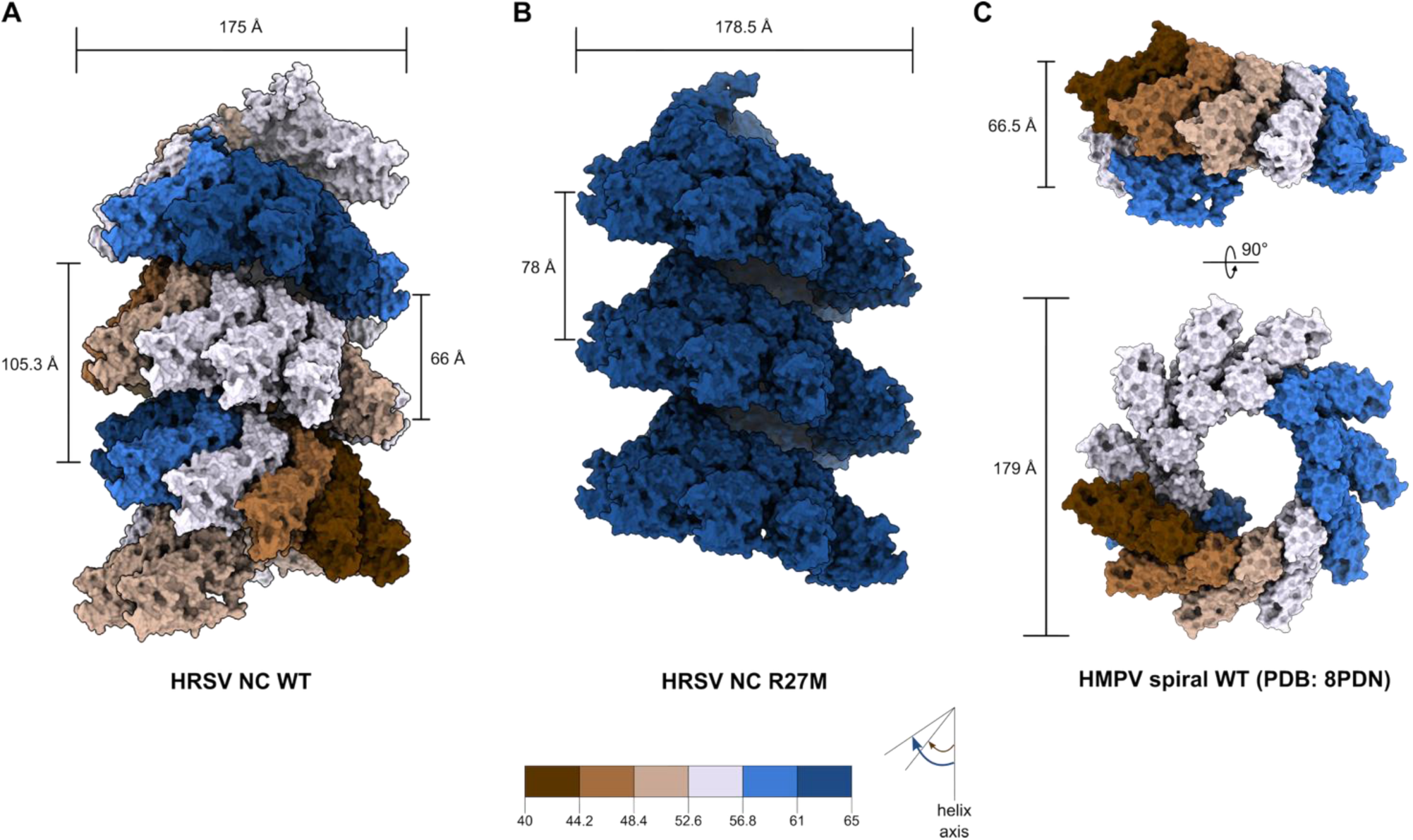
Helical symmetry comparison of HRSV WT NC, HRSV R27M NC and HPMV WT spiral. **(A)** Atomic model of the HRSV WT NC displayed as a surface. At the top, the NC diameter is indicated. On the left side, the size of one asymmetric unit (16 protomers) is indicated and on the right the helical-pseudo-pitch size is indicated. **(B)** Same as **A** but for HRSV R27M NC. At the top, the NC diameter is indicated. On the left side, the helical-pitch size is indicated. **(C)** Same as **A** but for HMPV WT spiral (PDB: 8PDN). On the left-top side, the helical-pseudo-pitch size is indicated. On the left-bottom side, the diameter of the spiral is indicated. Protomers are coloured depending on their axial tilt, the colour code is indicated at the bottom of the figure.

**Figure S5:**
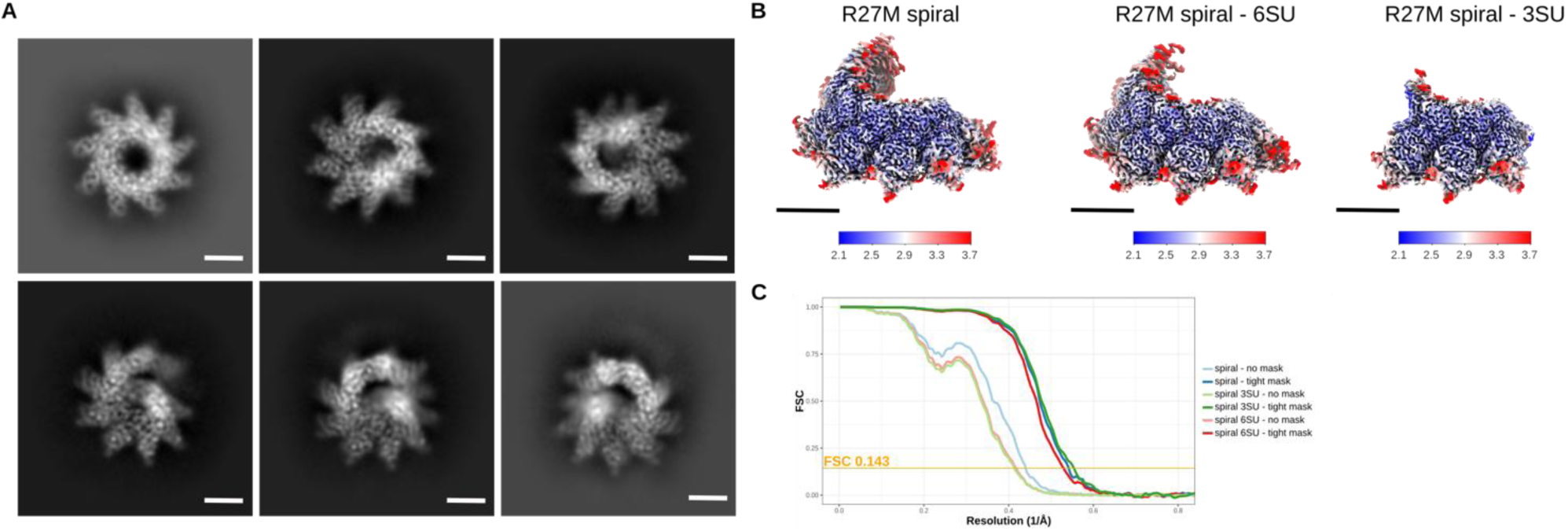
Additional information on cryo-EM analysis of R27M NC spirals. **(A)** Representative 2D class averages (top and tilted views). Scale bars, 50 Å. **(B)** Cryo-EM maps of the spiral and its six- and three-protomer subsections filtered and coloured by local resolution (in Å). Contour levels used in ChimeraX to generate the full views are 0.0925 (spiral), 0.08 (6SU subsection) and 0.095 (3SU subsection). Scale bars 50 Å. **(C)** FSC curves. Orange line represents FSC=0.143.

**Figure S6:**
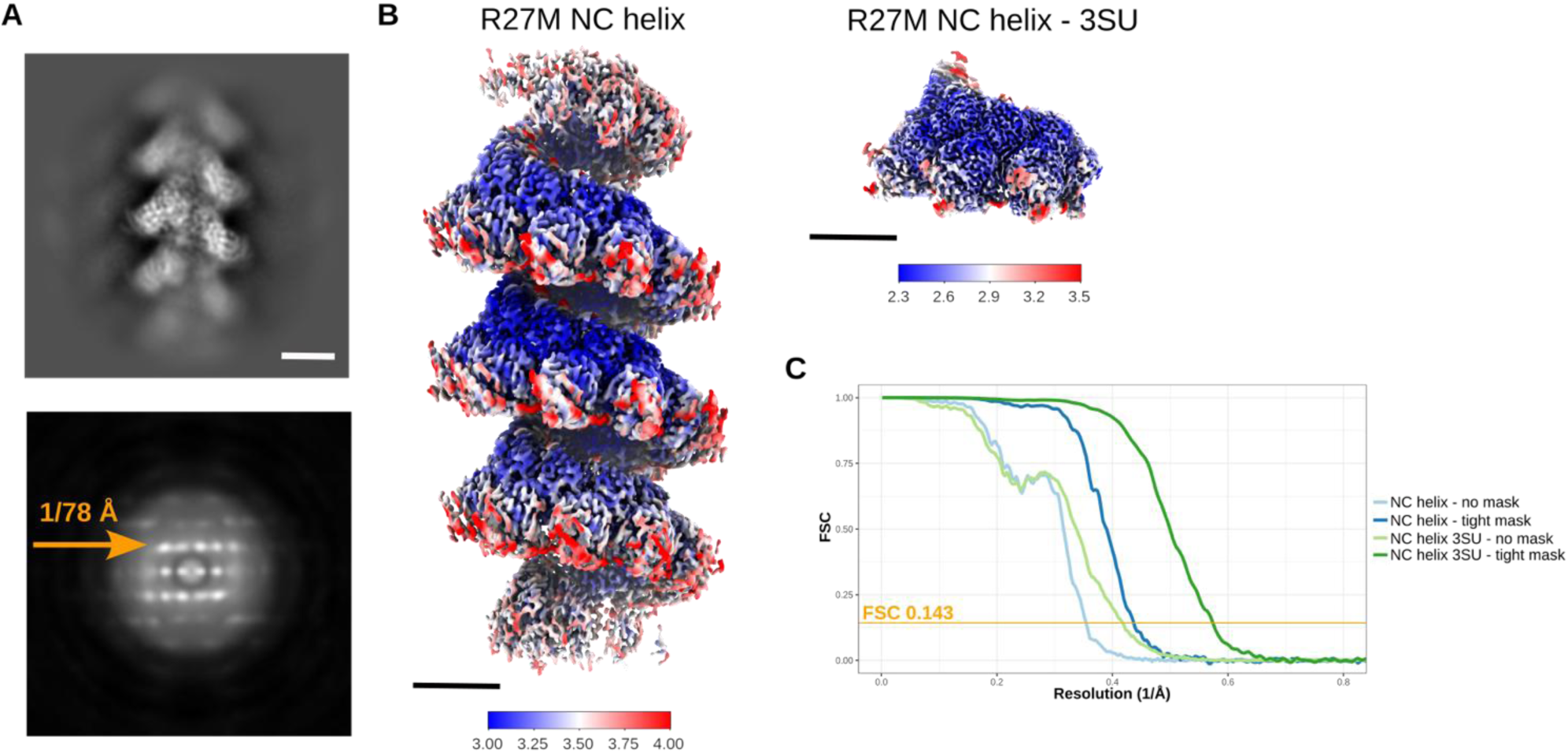
Additional information on cryo-EM analysis of R27M NC helices. **(A)** Representative 2D class average (top) of the R27M NC helix and an average of the PS of the particles in this class (bottom), showing a layer-line with the maximum close to the meridian at 1/78 Å. **(B)** Cryo-EM maps of the R27M NC helix and its three-protomer subsection filtered and coloured by local resolution (in Å). Contour levels used in ChimeraX to generate the full views are 0.055 and 0.10, respectively. Scale bars 50 Å. **(C)** FSC curves. Orange line represents FSC=0.143.

**Figure S7.**
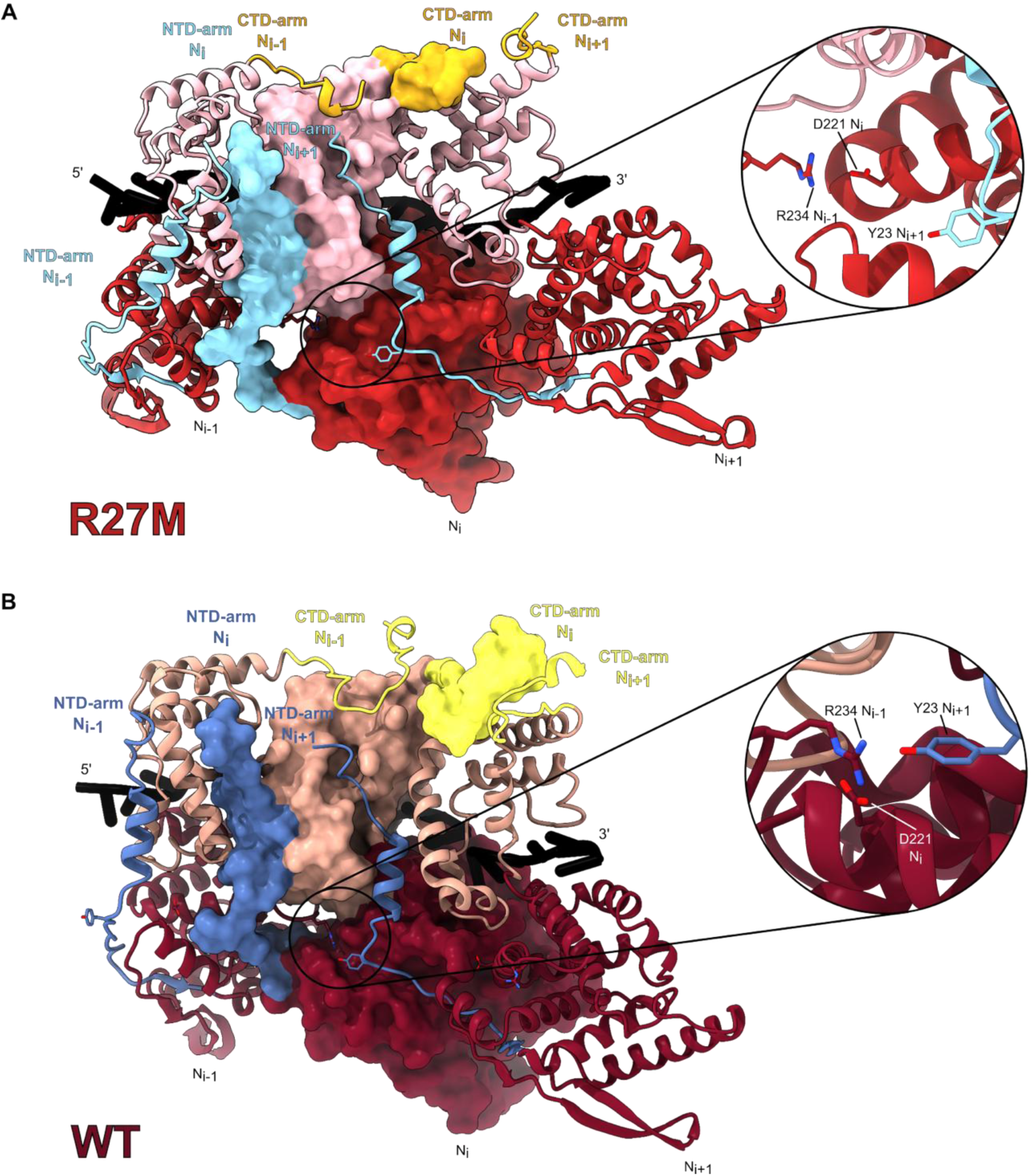
Lateral interactions (N-hole) between N protomers in HRSV R27M NC and HRSV WT N_10_ double rings. **(A)** Atomic models of three consecutive HRSV R27M NC protomers are shown, as ribbons on the edge and as surface for the middle one. The close-up of the N-hole shows the absence of tripartite interaction between Y23 and D221-R234. **(B)** Same as **A** but for the three protomers from HRSV WT N_10_ double rings (PDB: 8OOU). The close-up of the N-hole shows the tripartite interaction between Y23-D221-R234. The NTD-arms, NTD, CTD and CTD-arms are coloured as in Figures 1, 5 and 6.

**Supplementary Movie 1. Morphing of the WT helical nucleocapsid to the R27M helical nucleocapsid.** Atomic models of the HRSV WT helical subsection (PDB: 8OP1) and of the HRSV R27M three-protomer subsection helical nucleocapsid (the middle protomer of each atomic model is used for the alignment). Only the three-middle protomers of the HRSV WT helical subsection have been kept for the morphing. The NTD-arms, NTD, CTD and CTD-arms and RNAs are coloured as in Figures 1, 5 and 6.

